# Pump-Rest-Leak-Repeat: regulation of the mammalian-brain V-ATPase via ultra-slow mode-switching

**DOI:** 10.1101/2022.10.06.511076

**Authors:** Eleftherios Kosmidis, Christopher G. Shuttle, Julia Preobraschenski, Marcelo Ganzella, Peter J. Johnson, Salome Veshaguri, Mads P. Møller, Orestis Marantos, Jesper L. Pedersen, Reinhard Jahn, Dimitrios Stamou

## Abstract

Vacuolar-type adenosine triphosphatases (V-ATPases)^1–3^ are electrogenic rotary mechanoenzymes structurally related to F-type ATP synthases^4,5^. They hydrolyze ATP to establish electrochemical proton gradients for a plethora of cellular processes^1,3^. In neurons, the loading of all neurotransmitters into synaptic vesicles is energized by ~1 V-ATPase molecule per synaptic vesicle^6,7^. To shed light into this bona fide single-molecule biological process, we investigated electrogenic proton pumping by single mammalian-brain V-ATPases, using individual synaptic vesicles fused with immobilized liposomes. We show V-ATPases do not pump continuously in time, as hypothesized by observing the rotation of bacterial homologs^8^ and assuming strict ATP/proton coupling. Instead, they stochastically switch between three novel ultra-long-lived proton-pumping, inactive, and proton-leaky modes. Upending conventional wisdom, direct observation of pumping revealed that physiologically relevant concentrations of ATP do not regulate the intrinsic pumping rate. Instead, ATP regulates V-ATPase activity via the switching probability of the proton-pumping mode. In contrast, electrochemical proton gradients regulate the pumping rate and the switching of the pumping and inactive modes. This work reveals and emphasises the mechanistic and biological importance of mode-switching in protein regulation.

## Main Text

Vacuolar-type adenosine triphosphatases (V-ATPases) are a ubiquitous family of electrogenic ATP-driven rotary mechanoenzymes structurally related to F-type ATP synthases^4,5^. They use the energy released during ATP hydrolysis to pump protons across cellular membranes and are essential for acidification of intracellular organelles and for acid secretion^1,3^. Accordingly, their function is important for a plethora of diverse cellular, physiological and pathological processes including membrane trafficking, signaling, kidney function and cancer metastasis^3^.

In neurons, arguably the most important role of the V-ATPase is energizing the loading of all neurotransmitters in synaptic vesicles (SVs)^9–11^. In SVs the V-ATPase hydrolyzes ATP in the outward-facing soluble V_1_ region and converts the released chemical energy into rotation of the transmembrane V_0_ region that then translocates protons into the SV lumen through a coupling mechanism that is not well understood^9,12^ (Fig. 1a). The electrochemical proton gradient established by the V-ATPase subsequently energizes the secondary transport of all neurotransmitters into SVs^9,11,13^.

**Fig. 1.**
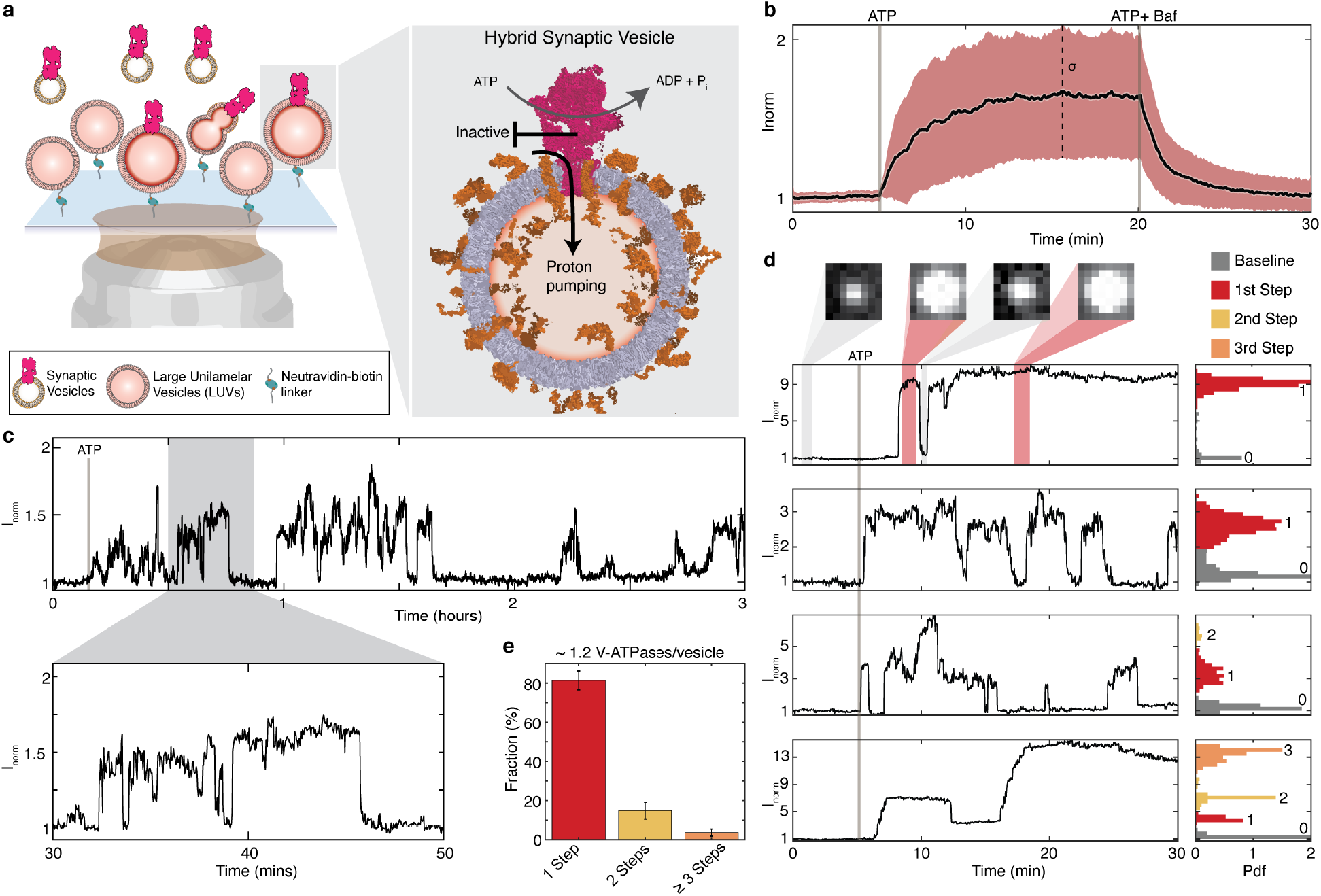
Ultra-long measurements of proton-pumping into single hybrid synaptic vesicles reveal mode-switching of individual mammalian brain V-ATPases. **a**, Illustration of assay for single transporter activity recordings. Proteoliposomes, carrying the pH sensitive fluorophore pHrodo, syntaxin1 and SNAP25, are immobilized on a glass slide. Synaptic vesicles containing mostly one copy of the V-ATPase are added in solution and fuse with the proteoliposomes, producing hybrid synaptic vesicles (SV_h_). Upon ATP hydrolysis the V-ATPase pumps protons into the lumen of SV_h_. **b**, Ensemble average of ~ 450 single SV_h_ acidification kinetics, data not corrected for photobleaching. Proton pumping and vesicle acidification were initiated by addition of ATP and inhibited by 200 nM bafilomycin. Black line is the mean, while the red-shaded area corresponds to one standard deviation. **c**, Ultra-long-term acidification kinetics of a typical single SV_h_ reveal the V-ATPase stochastically switches between long-lived proton-pumping and inactive modes. All kinetics have been processed by a Chung-Kennedy filter^24^ unless stated otherwise. **d**, Top two traces show one acidification plateau corresponding to one active V-ATPase per SV_h_. Bottom two traces are typical examples of SV_h_ traces with respectively two, three acidification plateaus corresponding to two, three active V-ATPases. **e**, Number of active V-ATPases per SV_h_ based on step-counting acidification plateaus. *n* = 8, where hereafter *n* is the number of independent experiments.

Despite the absolutely essential role of V-ATPases in loading SVs with neurotransmitters, each SV contains only one or two copies of the V-ATPase^6,7^. Thus, any putative single-molecule-based stochasticity in the function of the V-ATPase may be expected to bear a disproportionally high impact on SV loading and neuronal communication. To shed more light into this bona fide single-molecule biological process, we monitored here for the first time, to the best of our knowledge, electrogenic proton pumping by the V-ATPase at the single molecule level. Our measurements revealed three hitherto unknown ultra-long-lived modes that underlie the function of V-ATPases: (i) proton-pumping, (ii) inactive and (iii) proton-leaky mode. The regulation of V-ATPases by ATP and electrochemical gradients is based on switching between these modes, emphasising their mechanistic and biological importance.

### Isolating single mammalian-brain V-ATPases in single hybrid synaptic vesicles

To preserve the proton pumping activity of the mammalian brain V-ATPase, which is not as stable as its bacterial homologs^14^, we avoided detergent solubilization protocols and instead isolated the endogenous V-ATPase in intact SVs purified from rat brain^6,15^. Proton pumping of the V-ATPase could then in principle be observed by measuring the luminal pH of SVs using genetically encoded pH sensors^11,16^. However, their low expression levels (~4 copies per SV)^11^ have historically been limiting the signal/noise and duration of such recordings. Here, to observe the activity of the V-ATPase for periods of up to three hours we incorporated hundreds of photostable synthetic pH indicators (DOPE-pHrodo) per SV, using a well-tested protocol of SNARE-mediated vesicle fusion^17^.

As illustrated in Fig. 1a, we first tethered on a glass slide large unilamellar vesicles that contained DOPE-pHrodo and the SNARE acceptor complex, syntaxin 1 and SNAP 25 (LUVs). Subsequently, we injected SVs under conditions that we have previously demonstrated allow the fusion of predominantly one SV per LUV^18^ (Fig. 1a, Extended Data Fig. 1, 2). The outcome was immobilized hybrid synaptic vesicles (SV_h_) containing DOPE-pHrodo and the entire SV proteome, including the endogenous V-ATPase (Fig. 1a). As we and others have shown, the low surface density allowed observation of single vesicles^11,19^ while the controlled tethering conditions preserved membrane permeability to ions^19,20^, vesicle sphericity^21^ and the function of transmembrane proteins^19,22,23^.

### Ultra-long measurements of proton-pumping by single V-ATPases reveal mode-switching

To activate the V-ATPase and initiate proton pumping in the lumen of tethered SV_h_ we added ATP. Indeed, the ensemble average of many single SV_h_ traces confirms the expected ATP-specific decrease in luminal *pH*, shown as an increase in DOPE-pHrodo fluorescence in Fig. 1b (t = 5 min). The plateau reached after ~15 min reflects a dynamic equilibrium between protons being actively pumped into the lumen and protons passively permeating through the membrane down the concentration gradient^19,25^. Addition of the V-ATPase specific inhibitor bafilomycin (Fig. 1b, t = 20 min) blocked proton pumping and collapsed the proton gradient. In agreement with proteomic data^6^, this demonstrates that the V-ATPase is the only proton-pumping ATPase present in SV and is solely responsible for the luminal acidification observed in our experiment.

We next tracked the acidification of SV_h_ at the single particle level over periods of up to 3 hours comprising millions of proton turnovers^8^. Fig. 1c, Extended Data Fig. 3, show typical single SV_h_ traces. Strikingly, single SV_h_ kinetics revealed reversible stochastic pauses in the acidification that remained hidden in the ensemble average trace. In previous work with a P-type ATPase, we demonstrated that such pauses can be attributed directly to stochastic transitions of the transporter to long-lived inactive functional modes^19^. To confirm that the acidification pauses we observe here are indeed due to switching-off of the V-ATPase and not due to stochastic switching-on of a large leakage current either from one of the other proteins present in the SV_h_ or from the membrane itself, we performed the following control experiment. We imposed a pH gradient in the absence of ATP by exchanging the outside buffer and increasing the external pH by ~ 1 unit. This pH gradient resulted in an efflux of protons that we monitored in real time (Extended Data Fig. 4, Supplementary Methods). Efflux kinetics followed a continuous monotonic decay where no stochastic steps were observed. This finding confirms that when the V-ATPase is inactive there is no other proton-permeable pathway in the SV_h_ that can stochastically switch on/off. Thus, the acidification pauses can be directly attributed to the V-ATPase reversibly switching between a long-lived proton-pumping mode and an inactive mode.

A ‘mode’ comprises a plethora of ‘conformations’ (e.g. the continuous rotation taking place in the proton-pumping mode samples many discrete conformations^2,26^) which can be sampled millions of times before reversibly switching to another mode. The structure(s) underlying the inactive mode we observed here are not known at present. It is unlikely the inactive mode is due to the glucose-mediated dissociation/solubilization of the entire V_1_ region reported in yeast (because such dissociation would not be reversible at sub-nanomolar concentrations where we performed our experiments, see Supplementary Discussion) but the other apparently autoinhibited structures are plausible candidates^1^.

The ability to resolve stochastic mode-switching suggests the existence of a small number of single V-ATPase molecules per SV_h_, in agreement to proteomic data^6^. As we have shown previously, the presence of more than one transporter per vesicle leads to multiple steps/plateaus in the acidification traces^19^, similarly to having more than one ion channel in a membrane patch during a patch clamp recording. Here we could confirm this also for the V-ATPase (Fig. 1d). Step-analysis revealed that 81 % ± 5 % of the active SV_h_ showed a single acidification plateau indicative of a single active V-ATPase, whereas 15 % ± 5 % showed two plateaus, and 4 % ± 2 % three or more plateaus (Fig. 1e). Apart from directly reporting the distribution of active V-ATPase per SV, our results also show that on average there are 1.2 ± 0.1 active V-ATPases per SV. This is in excellent agreement with the ensemble average value of 1.4 measured by mass spectrometry^6^ and the average value of 1.27 ± 0.19 measured using fluorescent antibody labelling and single SV analysis^7^. These comparisons also suggest that ~85-95 % of all V-ATPases are active. We restricted all subsequent analysis to SV_h_ containing a single active V-ATPase.

### Reversible switching between proton-pumping, inactive and proton-leaky modes

To quantify the dynamics of the transitions between modes we developed a stochastic model to identify mode-switching events and calculate maximum a posteriori probability estimates of the signal (Fig. 2b, Extended Data Fig. 5, Supplementary Discussion). Utilizing the outputs of the stochastic model, we performed a dwell time analysis. Dwell time histograms were fitted well with single exponentials suggesting a single stochastic process underlies our observations (Fig. 2c, d). However, we cannot exclude the existence of other (faster or slower) modes outside the temporal resolution of our experiments. The characteristic dwell time of the pump in the proton-pumping mode was *τ_on_* = 174 s ± 9 s (Fig. 2c, Extended Data Fig 6b, top row), while pumping was interrupted by intervals of inactivity, *τ_off_* = 122 s ± 6 s (Fig. 2c, Extended Data Fig. 6c, top row). Thus, the probability the V-ATPase is actively pumping protons can be calculated to be *P_on_* = 0.6 ± 0.1. The fact the pump spends nearly half of the time switched off highlights the importance that modes outside the canonical pumping cycle bear for the function of V-ATPases. Ensemble average experiments that assume the V-ATPase to pump continuously over time would underestimate pumping rates by ~170 %.

**Fig. 2.**
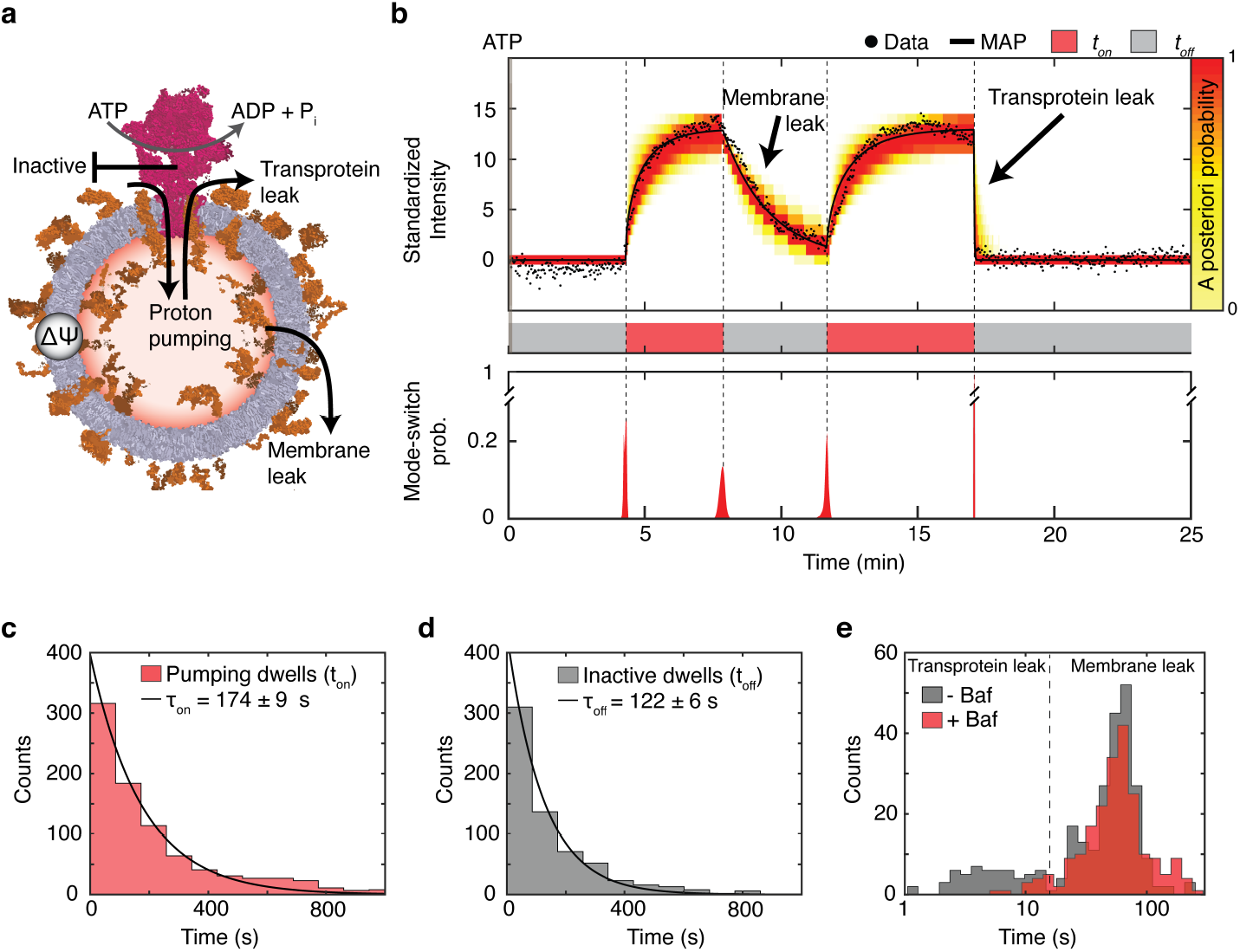
Dynamics of switching between the proton-pumping, inactive and proton-leaky modes. **a**, Illustration of different pathways for influx and efflux of protons in SV_h_. Proton pumping establishes a *pH* gradient and a membrane potential (*ΔΨ*). **b**, Analysis of stochastic fluctuations. Mode-switching events are detected using the a posteriori distributions (red) and accepted when certainty is > 99%. Stochastic filtering techniques are then used on the segmented data to find the parameters with the highest likelihood given the exponential model. The heat map represents the probability of where the signal is likely to be, given the observed data and the exponential model. The maximum a posteriori probability of the signal is seen here in black. **c**, **d**, Dwell time histograms for proton-pumping and inactive modes. Histograms were fit with single exponentials. *n* = *8*. **e**, Histogram of characteristic leakage lifetimes from the exponential fits shown in panel B. In the absence of bafilomycin, a fast and a slow component are observed with average leakage lifetimes of 6 s and 61 s. Upon addition of bafilomycin the fast leakage pathway is blocked and can be thus attributed to the activated V-ATPase. *n* = 15.

To investigate further the mechanisms of proton efflux after the V-ATPase is switched-off, we quantified the efflux kinetics by fitting them with a single exponential function. The histogram of characteristic efflux lifetimes in the presence of ATP revealed two separate peaks at 6 s ± 1 s and 61 *s* ± 1.0 s (Fig. 2e), demonstrating the existence of two distinct proton-leakage pathways in SV_h_ when the V-ATPase is active. We were able to attribute the larger lifetime (slow leakage) to passive leakage through the membrane because it coincided with the efflux lifetime in LUVs that did not contain any transporters. In agreement with this conclusion, SV_h_ leakage exhibited only passive leakage through the membrane when the V-ATPase was either inactive (- ATP) or specifically inhibited (+ bafilomycin) (see respectively Extended Data Fig. 4i, j, Fig. 2e). Thus, smaller leakage lifetimes (fast leakage) are specific to active V-ATPases. We considered the possibility leakage is due to reverse pumping occurring during ATP synthesis however the low concentration of adenosine diphosphate (ADP) in our samples due to autohydrolysis of ATP (~ nM) makes this highly unlikely.

We thus conclude that the active V-ATPase can exit the cycle of conformational states involved in pumping and switch to modes that can be either inactive or proton-leaking (Fig. 2a). Furthermore, the transition from the inactive to the leaky mode appears to be forbidden since the leaky mode is not observed when the V-ATPase is inactive (- ATP or + bafilomycin). Notably, the leakage mode is temporally distinct from the pumping mode. Thus, though the leakage mode reduces the ensemble average (and time average) pumping rates and ATP/net-proton-transport stoichiometry, it does not affect the pumping rates or the stoichiometry of the pumping mode. The leakage mode is thus conceptually and mechanistically distinct from substrate slippage hypothesised to occur stochastically in the thermodynamic pumping cycles^12,27^.

### Electrochemical proton gradients regulate the pumping rate and the lifetimes of the pumping and inactive modes

Though the biological purpose of the V-ATPase is to establish electrochemical proton gradients, little is known about gradient feedback on regulation^1,3,13^. Single-molecule studies of detergent reconstituted V-ATPase^8,28^ by definition cannot establish gradients, and ensemble average measurements (e.g. in fungal^27^ or plant^29^ vacuole) cannot distinguish the individual contributions of pumping, inactive and leaky modes to regulation. Here, we measured quantitively the proton gradients of individual SV_h_ via an in situ, single SV_h_-based, pH calibration, and deconvolved these effects (Extended Data Fig. 7 and Supplementary Methods).

We first established a proton-mediated electrochemical membrane potential by adding only ATP, while subsequently we added chloride for charge compensation (Fig. 3a, Extended Data Fig. 8). Chloride counterions enter the vesicle lumen through chloride/proton antiport^30^ and VGluT-mediated chloride influx^31,32^, and negate positive charge buildup thus releasing the electrical component of the gradient (*ΔΨ*) (i.e. *ΔΨ*_−*Cl*_ > *ΔΨ*_+*Cl*_ *≈ 0*, Extended Data Fig. 8a). In a separate control experiment we confirmed that chloride addition did not in itself introduce discernible acidification spikes (Extended Data Fig. 8b-d).

**Fig. 3.**
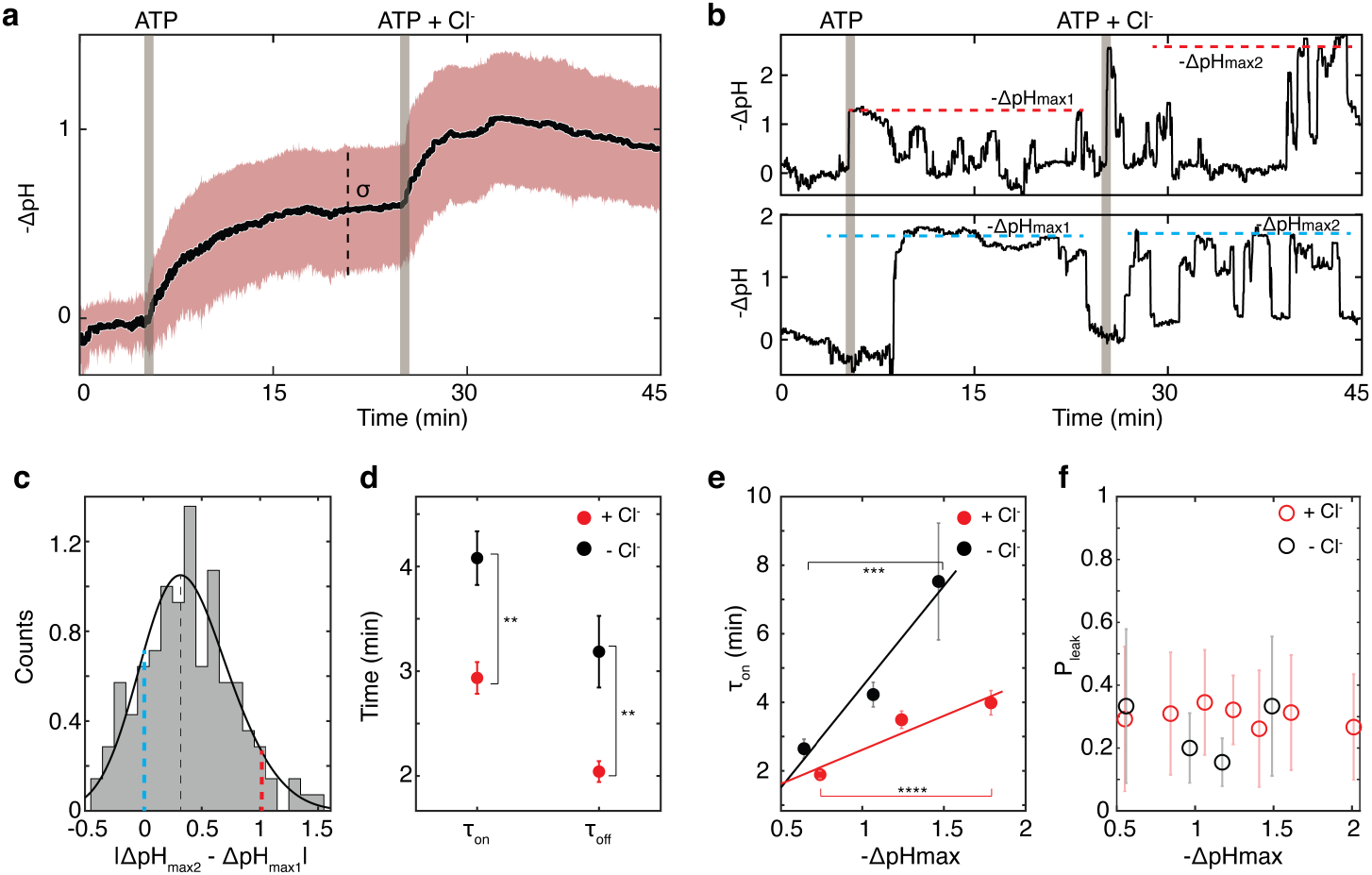
Mode-switching enables the regulation of the V-ATPase by electrochemical gradients. **a**, Ensemble average of acidification kinetics from ~ 220 single SV_h_, data not corrected for photobleaching. Proton pumping, and vesicle acidification was initiated by addition of ATP. Addition of choline chloride (+ Cl^−^) at 18 min collapsed the electrical component of the gradient (*ΔΨ*_−*Cl*_ > *ΔΨ*_+*Cl*_). Black line is the mean, while the red-shaded area corresponds to one standard deviation. −*ΔpH* is defined as the difference between the initial and the final *pH*. **b**, Representative examples of single SV_h_ acidification kinetics. Maximal acidification *(−ΔpH_max_*) is defined as the average value of pH plateaus achieved by the same single V-ATPase respectively under the two different conditions (± Cl^−^). **c**, Difference in maximum acidification. Colored lines correspond to the traces in panel (**b**) while black line to the mean. **d**, Characteristic dwell times for proton-pumping (*τ_on_*) and inactive (*τ_off_*) modes ± Cl^−^. Experiments used for dwell time determination were performed independently (Extended Data Fig. 6). A two-sided Mann-Whitney U-test produced P-values < 0.01 (**) for both modes. Total number of active traces used is *n* = 117 (− Cl^−^) and *n* = *467* (+ Cl^−^). Error bars correspond the error of single exponential fits on the total population of dwell times. **e**, proton-pumping dwell times (*τ_on_*) as a function of *−ΔpH_max_* (± Cl^−^). Data points and corresponding error bars have been produced by fitting single exponentials on equally sized subsets of the total population of dwell times. One-way ANOVA tests produced a *P*-value < 0.001 (+ Cl^−^) and *P-value* < 0.0001 (− Cl^−^). **f**, Transprotein leak probability as a function of *−ΔpH_max_*. Data points correspond to the medians of equally sized subsets of the total dataset. Error bars correspond to SD. Pearson correlation coefficients showed no linear correlation (*ρ* = −0.081, *P-value* = 0.25, − Cl^−^; *ρ* = 0.058, *P-value* = 0.09, + Cl^−^).

Ensemble average measurements confirmed that the release of *ΔΨ* regulates the V-ATPase and acidification on the average (Fig. 3a), in agreement with previous reports^16^. To deconvolve the pumping rates from the probability to be in the active mode we monitored acidification kinetics of single SV_h_ (Fig. 3b). This revealed, for the first time to our knowledge, that releasing *ΔΨ* indeed causes a bona fide increase in the (non-averaged) value of single SV_h_ maximal acidification (*ΔpH_max_*). The increase in maximal acidification (|*ΔpH_max2_* - *ΔpH_max1_*|) under the two conditions shows a broad distribution with a peak at ~ 0.3 pH units (Fig. 3c). We could thus verify that ~80 % of the single molecules increased their pumping rates when *ΔΨ* was released.

Notably, quantification of mode-switching dynamics also revealed potent regulation whereby the lifetimes of the proton-pumping mode (*τ_on_*) and of the inactive periods (*τ_off_*), were reduced respectively by 70 % and 60 % (Fig. 3d, Extended Data Fig. 6b, c, Extended Data Fig. 9). Lastly, these measurements suggest that the steady state value of *ΔpH_max_* = 1 which is quantified in ensemble average measurements^33^ as Fig. 3a, underestimates by ~180% the true maximal acidification values (Extended Data Fig. 8d) because it is weighted down by the periods of V-ATPase inactivity.

Next, we deconvolved the contribution of the electrical from the chemical part of the gradient to the regulation of *τ_on_*. *τ_on_* was positively correlated to increasing *ΔpH_max_* indicating the existence of a positive feedback loop intrinsic to the V-ATPase, whereby increasing the proton gradient from 0.75 to 1.75 enhances *τ_on_* by ~ 210 % (*P* < 0.0001, Fig. 3e, red dots). The positive correlation of *ΔpH* and *τ_on_* is not an artifact of slow build-up of chemical gradients, because kinetics of pH build-up are nearly an order of magnitude faster than the durations of on-cycle dwells (Extended Data Fig. 10). Increasing *ΔΨ*, further reinforces the positive feedback enhancing *τ_on_* by ~ 280 % (*P* < 0.0001, Fig. 3e, black dots). Thus, the pumping lifetime is positively correlated both to *ΔΨ* and *ΔpH*. In contrast, the probability of the V-ATPase to switch to the proton leaky mode was independent of *ΔpH* and *ΔΨ* (*ρ* = −0.081 and *ρ* = 0.058, Fig. 3f). A proton leaky mode has only been directly observed before for the eukaryotic H^+^-ATPase AHA2. However, in that case the probability to mode-switch was strongly regulated by proton gradients^19^.

Knowledge of *τ_on_* and *τ_off_* allowed us to use transition state theory to calculate the activation barriers for transiting between the proton-pumping and the inactive modes in the presence/absence of chloride (see Supplementary Discussion and Extended Data Fig. 11). The two activation barriers are very large (~ 86 kJ · mol^−1^) reflecting the stability of the conformations underlying the two long-lived modes that have dwell times of ~ 3 min (Fig. 3d). Chloride addition affects the forward and backward transitions nearly to the same extent, thus the overall effect on the probability to mode-switch is negligible (Extended Data Fig. 11a). This suggests that regulation by electrochemical gradients manifests mainly by modulating the transition barrier but not the ground state of the two modes (Extended Data Fig. 11b). The regulation of the activation barrier is small (~ 1 kJ mol^−1^, Extended Data Fig. 11), revealing subtle structural rearrangements which however have a dramatic influence on the dwell time of the pumping mode (up to ~ 280 %).

### ATP does not regulate the pumping rate, but only the probability to be in the proton-pumping mode

The availability of catalytic substrate (ATP) is of course essential for the proton pumping activity of the V-ATPase. Early on, single molecule rotation experiments directly observed how the rotation speed of the shaft (D subunit) was tuned by ATP concentration in a continuous Michaelis-Menten manner^8,34^. However, these experiments did not measure proton pumping and could thus not report how changes in the speed of ATP hydrolysis and rotation affect the proton current. Here, for the first time to our knowledge, we tested at the single molecule level the effect of ATP on proton pumping.

We explored a range of ATP concentrations from 10 μM to 1000 μM because the Michaelis-Menten constant (K_m_) of V-ATPases for ATP hydrolysis and rotation has been reported to be approximately 100 μM^34^. In agreement with previous reports, ensemble average acidification kinetics showed an ATP-dependent increase in the ensemble averaged proton pumping and *ΔpH* (Fig. 4a, Extended Data Fig. 12a, b)^16^. To deconvolve the lifetime from the rate of the proton pumping mode we investigated acidification kinetics of single SV_h_. and followed mode-switching over time at different concentrations of ATP (Fig. 4b).

**Fig. 4.**
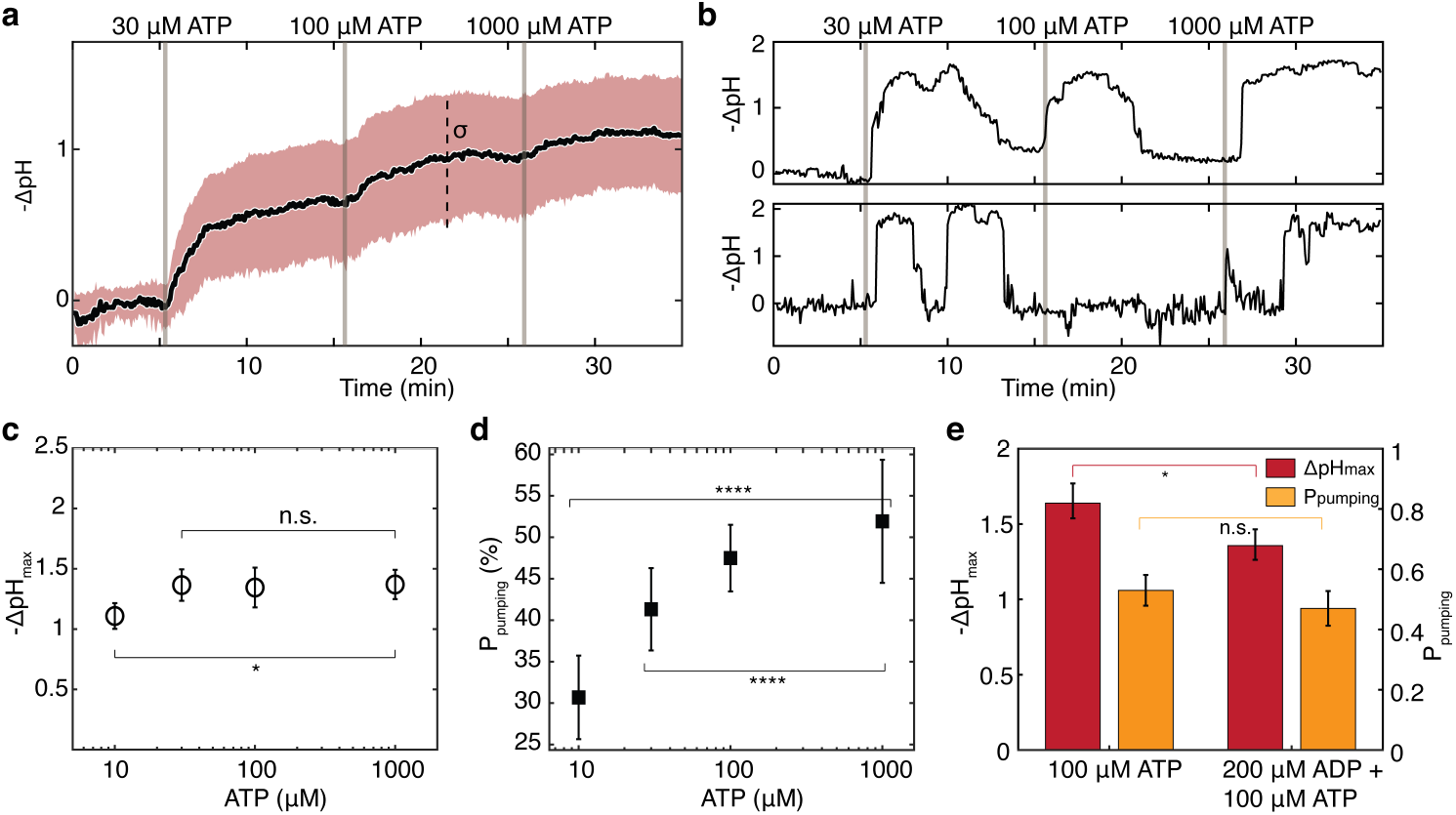
Regulation of functional modes by ATP and ADP. **a**, Ensemble average of 215 single hybrid vesicle acidification kinetics. Proton pumping and vesicle acidification was initiated by addition of ATP and Mg^2+^ in the constant presence of choline chloride throughout the duration of the experiment. Black line corresponds to the mean, while red-shaded area corresponds to the spread of one standard deviation in the single vesicle data. **b**, Characteristic single molecule data of proton pumping undergoing different ATP concentrations. It should be highlighted that the same single molecules are subjected to different concentrations of ATP. **c**, Pumping mode *ΔpH_max_* values as a function of ATP concentration. Data points correspond the means of *ΔpH_max_* between independent experiments. Error bars correspond to the SD between independent experiments. One-way ANOVA tests produced a *P-value > 0.05 (n.s.)* for [ATP] = 30 – 1000 μM and *P-value* < 0.05 (*) for [ATP] = 10 – 1000 μM. **d**, Probability that a molecule will be found in a pumping mode as a function of ATP. Data points correspond the means of *P_pumping_* between independent experiments. Error bars correspond to the SD between independent experiments. One-way ANOVA tests produced a *P-value* > 0.0001 (****) for [ATP] = 30 – 1000 μM and *P-value* < 0.0001 (****) for [ATP] = 10 – 1000 μM. Number of independent experiments is n = 4-7. The pumping efficiency of the brain V-ATPase does not depend on ATP concentration. However, the probability to be found on-cycle is drastically regulated by substrate concentration. **e**, *ΔpH_max_* of the pumping-mode showed a slight decrease in the presence of 200 μΜ ADP at a 2:1 ratio of ADP/ATP. Probability that the molecule is found in a pumping mode is not affected by the presence of ADP in the system. Bars points correspond to the mean and median of independent experiments for *ΔpH_max_* and *P_pumping_* respectively (*n* = 5). Error bars correspond to the SD between independent experiments. Two-sided Mann-Whitney U-test produced *P* < 0.05 and *P* > 0.05 for *ΔpH_max_* and *P_pumping_* respectively.

Paradoxically, the single-molecule experiments revealed that *ΔpH_max_* and thus the single-molecule pumping rate, remains constant between 30 μM and 1 mM ATP. (Fig. 4c, Extended Data Fig. 12c-e). Crucially, the increase observed in ensemble average proton pumping experiments originates solely to an ATP-depended regulation (up to 60%) of the probability (*P_pumping_*) that the V-ATPase is in the proton-pumping mode (Fig. 4d, Extended Data Fig. 13). Thus, this broad range of physiologically relevant ATP concentrations regulated solely mode-switching.

These experiments suggest that coupling of shaft rotation to proton shuttling across the membrane becomes the dominant rate limiting step in proton pumping well below the K_m_ and suggest further increase in the concentration of ATP results in futile cycles of hydrolysis and rotation. The reduction of *ΔpH_max_* at 10 μM ATP (Fig. 4c), suggests hydrolysis becomes rate limiting below 30 μM likely due to diffusion-limited binding of ATP to the V-ATPase^8^.

The molecular mechanism underlying ATP-mediated regulation of *P_pumping_* is not clear at present. We decided to test ADP/ATP competition because proton pumping by V-ATPases was reduced by half in yeast vacuoles upon addition of 5 mM ADP^27^ and ADP has been reported to induce inhibitory rotation pauses to isolated subdomains of V/F-type synthases^28,35,36^. We initiated acidification of SV_h_ with 100 μM ATP and after 20 minutes we competed with a two-fold excess of ADP in the presence of 5 mM phosphate (Extended Data Fig. 14a, b). Our measurements revealed competition by ADP caused a reduction (~ 20 %) of *ΔpH_max_* and the pumping rates, however, it had no discernible influence on *P_pumping_* (Fig. 4e, Extended Data Fig. 14c). This suggests that though inhibition by ADP affects the proton-pumping mode, it is not responsible for mode-switching between pumping and inactive modes.

To summarise, in eukaryotes there are two closely related rotary mechanoenzymes whose function is tightly coupled to proton transport across membranes: the F-type synthases that harness the energy of proton gradients to synthesise ATP, and the V-type ATPases that hydrolyse ATP to pump protons and establish proton gradients. They have both been investigated by pioneering single-molecule experiments that observed directly rotation of the central rotor γ-subunit within the isolated F_1_/V_1_ domain and validated the mechanism of rotation^37,8,38,39^, however, these experiments did not investigate proton transport. Two more recent interesting articles investigated proton transport by single bacterial F_0_F_1_-ATP synthases rotating backwards (i.e. hydrolysing ATP)^40,41^, however, the reported single-molecule pumping rates were averaged over periods of ~ 10 min masking any putative stochasticity.

Here we present, for the first time to our knowledge, rotary mechanoenzyme-mediated proton pumping under electrogenic conditions that allowed direct observation of stochastic fluctuations in pumping rates. Our measurements revealed that the native mammalian brain V-ATPase can exit the canonical proton-pumping mode and reversibly switch to physiologically relevant long-lived inactive and proton-leaky modes. This mode-switching introduces dramatic all/none fluctuations in the electrochemical gradient of SV_h_, and likely SVs, which should consequently lead to all/none fluctuations in the proton-coupled rates of secondary active neurotransmitter transport^11^. Such stochastic phenomena could help rationalise the reports of heterogenous neurotransmitter concentration in SVs^42^. In organelles comprising many transporters, mode-switching would not introduce as large fluctuations due to ensemble averaging, it would however directly determine the mean value of the gradient.

More broadly, mode-switching, also referred to as long-lived disorder in reaction rates, appears to be a general property of enzymes, however, its biological role has been unclear^43,44^. The observation that V-ATPase regulation by ATP and electrochemical gradients is based on mode-switching reveals a biological role of ultra-slow mode-switching/disorder in the regulation of protein function. Further elucidation of these phenomena will benefit greatly from direct, long-term, and where possible, simultaneous observation of conformational and functional dynamics, in conjunction with single-molecule guided determination of cryo-electron microscopy structures^19,23,45–49^.

## Methods

### Chemicals

Phospholipids and Cholesterol were purchased from Avanti Polar Lipids Inc. Chemicals for buffers, detergents and other reagents were purchased from Sigma-Aldrich, unless stated otherwise.

### The lipid-conjugated pH sensor PE-pHrodo

The lipid conjugated pH sensor PE-pHrodo was synthesized by linking DOPE to the pHrodo red ester reagent (Invitrogen, Eugene, OR, USA). Synthesis, purification and characterization of its properties as a function of pH are described in detail in^50^.

### Protein expression and purification

The stabilized Syntaxin/SNAP25 acceptor complex (ΔN complex) constructs consist of the pETDuet-1 vector carrying syntaxin-1A (183-288) and the C-terminal fragment of synaptobrevin 2 (49-96) and the pET28a vector carrying His_6_-tagged SNAP-25A^51,52^ and originate from *Rattus norvegicus*. The components of the deltaN complex, syntaxin-1A (183-288), synaptobrevin 2 (49-96) and SNAP-25A were co-expressed in *E.coli* BL21(DE3) and purified as previously described^51,52^. In brief, the cells were lysed, and the complex was extracted using 10 % Na-Cholate (w/v) in 4 M urea, 500 mM NaCl, 20 mM HEPES, pH 7.4 and 8 mM imidazole. The non-dissolved particles were separated by centrifugation and the cleared lysate was subjected to Ni^2+^-NTA affinity purification. Finally, the complex was further purified by anion exchange chromatography in presence of 2% CHAPS using the ÄKTA system (GE Healthcare), flash frozen and stored at −80 °C until use.

### Proteoliposome preparation and characterization

Proteoliposomes were prepared by mixing the detergent-solubilized ΔN complex together with lipids and subsequently removing the detergent by dialysis^53,54^. In detail, the synthetic lipid mix consisting of 1,2.dioleoyl-*sn*-glycero-3-phosphocholine (DOPC), 1,2-dioleoyl-*sn*-glycero-3-phospho-L-serine (DOPS) (both Avanti Polar Lipids), cholesterol (Chol) (from sheep wool, Avanti Polar Lipids), 1,2-dioleoyl-sn-glycero-3-phosphoethanolamine-N-(cap biotinyl) (18:1 Biotinyl Cap PE, Avanti Polar Lipids) and DOPE-pHrodo (see above) at a molar ratio of DOPC:DOPS:Chol:(18:1 Biotinyl Cap PE):DOPE-pHrodo 64.4:10:25:0.5:0.1 was formed by evaporating the organic solvent from the lipids and dissolving the dried film in 300 mM glycine, 2 mM MOPS-Tris, pH 7.3, 2 mM MgSO_4_, 5 % n-Octyl-beta-D-glucopyranosid to a concentration of 10 mg/ml lipids. The protein:lipid ratio (mol:mol) was adjusted to 1:650 and the final lipid concentration was adjusted to 4 mM. After mixing, the respective solutions were dialysed in Slide-A-Lyzer dialysis cassettes (2 kDa MWCO, Thermo Scientific) over night at 4 °C in 300 mM glycine, 2 mM MOPS-Tris, pH 7.3, and 2 mM MgSO_4_. The dialysis buffer was additionally supplemented with 2 g BioBeads (BioRad)/l buffer to adsorb detergent monomers. After dialysis, liposomes were aliquoted, flash frozen and stored at −80 °C until use. Prior to use, proteoliposomes were subjected to sequential extrusion through 800 nm and 200 nm size nucleopore polycarbonate membranes mounted on a mini extruder (Avanti Polar Lipids, Alabaster, AL).

### Synaptic vesicles from rat brain

Synaptic vesicles (SVs) were isolated from rat brain according to previous publications^6,55,56^. In brief, an SV-enriched fraction LP2 was prepared by differential centrifugation and subjected to a continuous sucrose density gradient centrifugation. The part between 0.04 and 0.4 M sucrose (colored) was collected and additionally separated by chromatography on controlled-pore glass beads (CPG). After the size exclusion chromatography, the SVs were pelleted by ultracentrifugation, resuspended in 320 mM sucrose and 5 mM HEPES, pH 7.4, flash frozen and stored at −80 °C until use. For more details see also Extended Data Fig. 1A. The protein concentration range varies from 1.5 mg/ml to 2.5 mg/ml.

Adult Wistar rats were purchased from Charles River Laboratories, or Janvier, and were kept until 5 to 6 weeks of age at a 12:12 hr light/dark cycle with food and water ad libitum. Certificate of approval for using animals was issued by the Landkreis Götitngen Office of Veterinary Affairs and Consumer Protection.

### Ensemble average ATPase assay

The activity of each preparation of SVs and pHrodo-labelled proteoliposomes was assessed by generating hybrid SVs as previously described^32,57^ and measuring chloride-dependent acidification of the hybrid SVs in a bulk assay using pHrodo as a pH sensitive probe. In brief, 60 μl pHrodo-labelled +ΔN liposomes were preincubated with 10 μl of SVs (c = 2 mg/ml) 45 min at room temperature. Subsequently, the acidification of the pHrodo-labelled hybrid SVs was monitored by measuring changes in fluorescence at 590 nm (slits: 3 nm for excitation at 560 nm and 5 nm for emission at 590 nm) in a fluorimeter (Fluoromax-2, Horiba) at 37 °C. Typically, 930 μl buffer (300 mM glycine, 30 mM choline chloride, 2 mM MOPS-Tris, pH 7.3, 2 mM MgSO_4_) were mixed in a cuvette with 70 μL hybrid SVs. The acidification reaction was started by adding 2 mM MgATP and stopped by adding 20 mM (NH_4_)_2_SO_4_. Representative traces are shown in Extended Data Fig. 2.

### Surface preparation and immobilization of +ΔN proteoliposomes

Surfaces were prepared according to previously published protocols^19,22^. Briefly, glass slides of thickness 0.17 ± 0.01 mm passed sequential sonication cycles with 2% (v/v) Helmanex III, MilliQ and subsequently rinsed with 99% (v/v) Ethanol and 99% (v/v) Methanol. Then, the slides were dried under nitrogen flow and plasma etched for three minutes to remove impurities. The glass slide is attached to Ibidi sticky slides VI 0.4. The surface is functionalized by first incubating a mixture of 100:1 PLL-g-PEG and PLL-g-PEG-biotin at 1mg/ml (SuSoS) for 30 minutes and then washed away with a 15 mM HEPES buffer. 0.1 mg/ml Neutravidin in 15 mM HEPES buffer is then incubated for 10 minutes and flushed out using the activity buffer (300 mM glycine, 2 mM MOPS, 2mM MgSO4, pH 7.1 at 23 °C. pH is adjusted with Tris). Once the flow chamber has equilibrated with the activity buffer, we exchange the buffer with the proteoliposome solution (proteoliposomes in activity buffer) and incubate until we reach a density of approximately 1000 immobilized liposomes in a 81.92 μm × 81.92 μm field of view (FOV). Subsequently, non-immobilized vesicles are flushed away.

### Hybrid synaptic vesicle formation

Purified synaptic vesicles are thawed, diluted (1:100) and incubated in the activity buffer for 5-10 minutes. Once spatially separated LUVs + ΔN (henceforth referred to as LUVs) are immobilized on the functionalized glass surface, SVs are injected into the flow chamber containing the proteoliposomes allowing for the fusion reaction to commence. Typically, incubation of SVs in the flow chamber lasted approximately 45 minutes. SV_h_ were prepared using the aforementioned method in all experiments apart from the ones shown in Fig. 1c and Fig. 2e, where the fusion reaction was performed in solution, in Eppendorf tubes, by mixing the populations of SVs and LUVs and allowing for 45-minute incubation. Then, SV_h_ were directly immobilized onto the glass surface.

### Image acquisition

All fluorescence microscopy was performed on a commercial Olympus total internal reflection fluorescence (TIRF) microscope (Olympus Europa SE & Co. KG, Hamburg, Germany). We used the EMCCD camera iXon 897 (Andor Technology, Belfast, UK). Excitation of fluorophores was achieved via an Olympus Cell* solid-state laser emitting at 532 nm. The microscope was equipped with an Olympus TIRF UApoN x100, 1.49 NA, immersion oil objective. The microscope was able to maintain focus for extended periods of time via the continuous function of the Zero Drift Correction module built into the microscope. An excitation bandpass filter 532/10 was used for fluorophore excitation. An emission filter with a bandpass 582.5/75 and a beam splitter bandpass 585/75 were used to block fluorophore excitation. Image acquisition software linked to the microscope and used throughout this article was for the most part Olympus xcellence rt and to a lesser extend Olympus cellSens.

Replicates of measurements for each condition were taken on the same sample measured repeatedly throughout the present manuscript.

### Reporting summary

Further information on research design is available in the Nature Research Reporting Summary linked to this paper.

## Data availability

The data that support the findings are available from the corresponding authors upon reasonable request.

## Code availability

Source code and demo for the Bayesian filtering algorithm which was used for the detection of events can be found in https://github.com/pete906/Bayesian_Filtering_Proton_Pump.git. Support on the use of the algorithm can be provided by the corresponding authors upon reasonable request.

## Acknowledgements

We would like to thank S.C. Blanchard and M. Grabe for inspiring discussions. H. Grubmüller, C. Kutzner and P.E. Gourdon for providing and helping with visualizations of the synaptic vesicle. This work was supported by the Novo Nordisk Foundation (Grant NNF17OC0028176), the Villum Foundation (Grants 17617 and 17646) and the Lundbeck Foundation (Grant R249-2017-1406 to E.K. and R250-2017-1175 to S.V.), and a grant from the European Research Council to RJ (SVNeuroTrans).

## Author contributions

D.S. conceived the strategy and was responsible for project management and supervision. E.K., J.P. and D.S. designed research with initial help from S.V. E.K. developed the single molecule assay and collected and analysed all data. C.S. was the principal software developer. J.P. prepared all biochemical samples, with help from M.G., under the supervision of R.J. P.J.J and J.L.P. developed the stochastic event detection model and estimations. M.P.M. established the pH calibration. O.M. simulated the spatial distribution of vesicles. D.S. and E.K. wrote the main text. E.K. prepared all figures and SI. All authors discussed the results and commented on the manuscript.

## Competing interests

The authors declare no competing interests.

**Supplementary information** is available for this paper

## Notes

### Competing Interest Statement

The authors have declared no competing interest.

## References

1. Vasanthakumar, T. & Rubinstein, J. L. Structure and Roles of V-type ATPases. Trends Biochem. Sci. 45, 295–307 (2020).

2. Ueno, H., Suzuki, K. & Murata, T. Structure and dynamics of rotary V1 motor. Cell. Mol. Life Sci. 75, 1789–1802 (2018).

3. Forgac, M. Vacuolar ATPases: rotary proton pumps in physiology and pathophysiology. Nat. Rev. Mol. Cell Biol. 8, 917–929 (2007).

4. Spikes, T. E., Montgomery, M. G. & Walker, J. E. Structure of the dimeric ATP synthase from bovine mitochondria. Proc. Natl. Acad. Sci. 117, 23519–23526 (2020).

5. Okuno, D., Iino, R. & Noji, H. Rotation and structure of FoF1-ATP synthase. J. Biochem. (Tokyo) 149, 655–664 (2011).

6. Takamori, S. et al. Molecular Anatomy of a Trafficking Organelle. Cell 127, 831–846 (2006).

7. Mutch, S. A. et al. Protein Quantification at the Single Vesicle Level Reveals That a Subset of Synaptic Vesicle Proteins Are Trafficked with High Precision. J. Neurosci. 31, 1461–1470 (2011).

8. Furuike, S. et al. Resolving stepping rotation in Thermus thermophilus H+-ATPase/synthase with an essentially drag-free probe. Nat. Commun. 2, 233 (2011).

9. Abbas, Y. M., Wu, D., Bueler, S. A., Robinson, C. V. & Rubinstein, J. L. Structure of V-ATPase from the mammalian brain. Science 367, 1240–1246 (2020).

10. Rost, B. R. et al. Optogenetic acidification of synaptic vesicles and lysosomes. Nat. Neurosci. 18, 1845–1852 (2015).

11. Farsi, Z. et al. Single-vesicle imaging reveals different transport mechanisms between glutamatergic and GABAergic vesicles. Science 351, 981–984 (2016).

12. Grabe, M., Wang, H. & Oster, G. The Mechanochemistry of V-ATPase Proton Pumps. Biophys. J. 78, 2798–2813 (2000).

13. Gowrisankaran, S. & Milosevic, I. Regulation of synaptic vesicle acidification at the neuronal synapse. IUBMB Life 72, 568–576 (2020).

14. Dilworth, M. V., Findlay, H. E. & Booth, P. J. Detergent-free purification and reconstitution of functional human serotonin transporter (SERT) using diisobutylene maleic acid (DIBMA) copolymer. Biochim. Biophys. Acta BBA - Biomembr. 1863, 183602 (2021).

15. Ahmed, S., Holt, M., Riedel, D. & Jahn, R. Small-scale isolation of synaptic vesicles from mammalian brain. Nat. Protoc. 8, 998–1009 (2013).

16. Budzinski, K. L., Zeigler, M., Fujimoto, B. S., Bajjalieh, S. M. & Chiu, D. T. Measurements of the Acidification Kinetics of Single SynaptopHluorin Vesicles. Biophys. J. 101, 1580–1589 (2011).

17. Hernandez, J. M. et al. Membrane Fusion Intermediates via Directional and Full Assembly of the SNARE Complex. Science 336, 1581–1584 (2012).

18. Castorph, S. et al. Synaptic vesicles studied by dynamic light scattering. Eur. Phys. J. E 34, 63 (2011).

19. Veshaguri, S. et al. Direct observation of proton pumping by a eukaryotic P-type ATPase. Science 351, 1469–1473 (2016).

20. Stamou, D., Duschl, C., Delamarche, E. & Vogel, H. Self-Assembled Microarrays of Attoliter Molecular Vessels. Angew. Chem. Int. Ed. 42, 5580–5583 (2003).

21. Bendix, P. M., Pedersen, M. S. & Stamou, D. Quantification of nano-scale intermembrane contact areas by using fluorescence resonance energy transfer. Proc. Natl. Acad. Sci. 106, 12341–12346 (2009).

22. Mathiasen, S. et al. Nanoscale high-content analysis using compositional heterogeneities of single proteoliposomes. Nat. Methods 11, 931–934 (2014).

23. Fitzgerald, G. A. et al. Quantifying secondary transport at single-molecule resolution. Nature 575, 528–534 (2019).

24. Chung, S. H. & Kennedy, R. A. Forward-backward non-linear filtering technique for extracting small biological signals from noise. J. Neurosci. Methods 40, 71–86 (1991).

25. Singh, A. et al. Protons in small spaces: Discrete simulations of vesicle acidification. PLOS Comput. Biol. 15, e1007539 (2019).

26. Zhao, J., Benlekbir, S. & Rubinstein, J. L. Electron cryomicroscopy observation of rotational states in a eukaryotic V-ATPase. Nature 521, 241–245 (2015).

27. Kettner, C., Bertl, A., Obermeyer, G., Slayman, C. & Bihler, H. Electrophysiological Analysis of the Yeast V-Type Proton Pump: Variable Coupling Ratio and Proton Shunt. Biophys. J. 85, 3730–3738 (2003).

28. Uner, N. E. et al. Single-molecule Analysis of Inhibitory Pausing States of V1-ATPase. J. Biol. Chem. 287, 28327–28335 (2012).

29. Davies, J. M., Hunt, I. & Sanders, D. Vacuolar H(+)-pumping ATPase variable transport coupling ratio controlled by pH. Proc. Natl. Acad. Sci. 91, 8547–8551 (1994).

30. Accardi, A. Structure and gating of CLC channels and exchangers: Structure and gating of CLC channels and exchangers. J. Physiol. 593, 4129–4138 (2015).

31. Schenck, S., Wojcik, S. M., Brose, N. & Takamori, S. A chloride conductance in VGLUT1 underlies maximal glutamate loading into synaptic vesicles. Nat. Neurosci. 12, 156–162 (2009).

32. Preobraschenski, J., Zander, J.-F., Suzuki, T., Ahnert-Hilger, G. & Jahn, R. Vesicular Glutamate Transporters Use Flexible Anion and Cation Binding Sites for Efficient Accumulation of Neurotransmitter. Neuron 84, 1287–1301 (2014).

33. Maycox, P. R., Deckwerth, T., Hell, J. W. & Jahn, R. Glutamate uptake by brain synaptic vesicles. Energy dependence of transport and functional reconstitution in proteoliposomes. J. Biol. Chem. 263, 15423–15428 (1988).

34. Minagawa, Y. et al. Basic Properties of Rotary Dynamics of the Molecular Motor Enterococcus hirae V1-ATPase. J. Biol. Chem. 288, 32700–32707 (2013).

35. Kishikawa, J., Nakanishi, A., Furuike, S., Tamakoshi, M. & Yokoyama, K. Molecular Basis of ADP Inhibition of Vacuolar (V)-type ATPase/Synthase. J. Biol. Chem. 289, 403–412 (2014).

36. Nakanishi, A., Kishikawa, J., Tamakoshi, M., Mitsuoka, K. & Yokoyama, K. Cryo EM structure of intact rotary H+-ATPase/synthase from Thermus thermophilus. Nat. Commun. 9, 89 (2018).

37. Yasuda, R., Noji, H., Yoshida, M., Kinosita, K. & Itoh, H. Resolution of distinct rotational substeps by submillisecond kinetic analysis of F1-ATPase. Nature 410, 898–904 (2001).

38. Noji, H., Yoshida, M. & Kinosita, K. Direct observation of the rotation of F1-ATPase. Nature 4 (1997).

39. Adachi, K. et al. Coupling of Rotation and Catalysis in F1-ATPase Revealed by Single-Molecule Imaging and Manipulation. Cell 130, 309–321 (2007).

40. Watanabe, R. et al. Arrayed lipid bilayer chambers allow single-molecule analysis of membrane transporter activity. Nat. Commun. 5, 4519 (2014).

41. Soga, N. et al. Monodisperse Liposomes with Femtoliter Volume Enable Quantitative Digital Bioassays of Membrane Transporters and Cell-Free Gene Expression. ACS Nano 14, 11700–11711 (2020).

42. Phan, N. T. N., Li, X. & Ewing, A. G. Measuring synaptic vesicles using cellular electrochemistry and nanoscale molecular imaging. Nat. Rev. Chem. 1, 0048 (2017).

43. Lu, H. P., Xun, L. & Xie, X. S. Single-Molecule Enzymatic Dynamics. Science 282, 1877 (1998).

44. Ciftci, D. et al. Single-molecule transport kinetics of a glutamate transporter homolog shows static disorder. Sci. Adv. 6, eaaz1949 (2020).

45. Akyuz, N., Altman, R. B., Blanchard, S. C. & Boudker, O. Transport dynamics in a glutamate transporter homologue. Nature 502, 114–118 (2013).

46. Erkens, G. B., Hänelt, I., Goudsmits, J. M. H., Slotboom, D. J. & van Oijen, A. M. Unsynchronised subunit motion in single trimeric sodium-coupled aspartate transporters. Nature 502, 119–123 (2013).

47. Akyuz, N. et al. Transport domain unlocking sets the uptake rate of an aspartate transporter. Nature 518, 68–73 (2015).

48. Dyla, M. et al. Dynamics of P-type ATPase transport revealed by single-molecule FRET. Nature 551, 346–351 (2017).

49. Rundlet, E. J. et al. Structural basis of early translocation events on the ribosome. Nature 595, 741–745 (2021).

50. Kemmer, G. C. et al. Lipid-conjugated fluorescent pH sensors for monitoring pH changes in reconstituted membrane systems. The Analyst 140, 6313–6320 (2015).

51. Pobbati, A. V. N- to C-Terminal SNARE Complex Assembly Promotes Rapid Membrane Fusion. Science 313, 673–676 (2006).

52. Stein, A., Radhakrishnan, A., Riedel, D., Fasshauer, D. & Jahn, R. Synaptotagmin activates membrane fusion through a Ca2+-dependent trans interaction with phospholipids. Nat. Struct. Mol. Biol. 14, 904–911 (2007).

53. Rigaud, J.-L. & Lévy, D. Reconstitution of Membrane Proteins into Liposomes. in Methods in Enzymology vol. 372 65–86 (Elsevier, 2003).

54. Rigaud, J.-L., Pitard, B. & Levy, D. Reconstitution of membrane proteins into liposomes: application to energy-transducing membrane proteins. Biochim. Biophys. Acta BBA - Bioenerg. 1231, 223–246 (1995).

55. Huttner, W. B., Schiebler, W., Greengard, P. & De Camilli, P. Synapsin I (protein I), a nerve terminal-specific phosphoprotein. III. Its association with synaptic vesicles studied in a highly purified synaptic vesicle preparation. J. Cell Biol. 96, 1374–1388 (1983).

56. Nagy, A., Baker, R. R., Morris, S. J. & Whittaker, V. P. The preparation and characterization of synaptic vesicles of high purity. Brain Res. 109, 285–309 (1976).

57. Preobraschenski, J. et al. Dual and Direction-Selective Mechanisms of Phosphate Transport by the Vesicular Glutamate Transporter. Cell Rep. 23, 535–545 (2018).

