## Supplementary Information for "Pump-Rest-Leak-Repeat: regulation of the mammalian-brain V-ATPase via ultra-slow mode-switching"

### Supplementary discussion

#### Stochastic model description

In the following the 'signal' will refer to the increased intensities of light in the experiment due to the V-ATPase being active. The signal depends on change-point times $\boldsymbol{T} = \{T_1, T_2, \ldots, T_n\}$ which are the times of mode-switching events at which the V-ATPase switches from an on-cycle state to an off-cycle state or visa versa. These states alternate, such that $T_i$ represents the point at which the proton pump switches to an proton-pumping mode if $i$ is odd or an inactive mode if $i$ is even.

In the proton-pumping mode, protons are pumped inside the vesicle, and so the fluorescence increases. This is modelled as an exponential increase up to a dynamic equilibrium of the system at which point the influx of protons due to the V-ATPase is equal to the passive efflux through the membrane. This equilibrium level we call $\alpha$ and the exponential rate as $\beta$. Then when the V-ATPase switches to an inactive-mode we model the fluorescence decreasing exponentially back to the baseline. The rate of this exponential decay we call $\gamma$. These exponential increases and decays from a general starting point K, between the baseline and equilibrium level $\alpha$, can then be represented as follows $\mu_1(t; K) = (K - \alpha)\exp(-\beta t) + \alpha$ and $\mu_2(t; K) = K\exp(-\gamma t)$. As the signal is switching state with every mode-switching event the general formula/model for the signal is

$$\mu(t) = \begin{cases} \mu_2(t; 0) & if \quad 0 \leq t < T_1 \\ \mu_1(t - T_1; \mu(T_1)) & if \quad T_1 \leq t < T_2 \\ \mu_2(t - T_2; \mu(T_2)) & if \quad T_2 \leq t < T_3 \\ \mu_1(t - T_3; \mu(T_3)) & if \quad T_3 \leq t < T_4 \\ \vdots & \vdots \end{cases}$$

noting that $\mu_2(t; 0) = 0$.

In this setting, we allow the exponential rates to be different from those seen previously (but fixed between mode-switching events), so for example we will denote $\beta_1$ and $\gamma_1$ to be the exponential rates of the first proton-pumping event and first inactive event respectively. However, we will assume the equilibrium level for the cell $\alpha$ to be the same for any point in time. For ease of notation, we will group these parameters together as $\boldsymbol{\nu} = \{\alpha, \beta_1, \gamma_1, \ldots, \beta_n, \gamma_n\}$. Letting the time between mode-switching events be defined as $\boldsymbol{\theta} = \{\theta_1, \theta_2, \ldots, \theta_n\} = \{T_1, T_2 - T_1, \ldots, T_n - T_{n-1}\}$, and note that the signal's dependance on $\boldsymbol{T}$ can be replaced by $\boldsymbol{\theta}$ as one set can be constructed from the other. We shall write the general formula for the signal as $\mu(t, \boldsymbol{\theta}, \boldsymbol{\nu})$.

Each signal depends uniquely on the equilibrium level for the vesicle, exponential rates in the model and also the times between mode-switching events which are not known a priori. These unknows are modelled as random variables with given unbiased prior distributions. In this Bayesian setting, it is assumed that all the random variables are mutually independent and the distributions are given as follows; the time between change-points is exponentially distributed (highest entropy on the positive real numbers); the equilibrium level for the cell α is uniformly distributed (highest entropy over a given interval); the distribution for the exponential rates $\beta$ and $\gamma$ are a uniform distribution over an approximately even spread for the slopes of the exponential functions $\mu_i$. Using unbiased prior distributions ensures that these priors have the least impact on the a posteriori estimates that are calculated (see next section).

**Stochastic filtering**

This section looks to give a brief overview of how techniques from Bayesian stochastic filtering are used in the context of this paper. For an introduction to stochastic filtering techniques see[1]. Bayesian stochastic filtering is a method of filtering out the unwanted noise leaving an estimate of the signal (given the model described above). The method takes the a priori opinion of the signal's change-points and parameters in the form of the unbiased prior distributions and provides an updated opinion given the data, known as the a posteriori (joint) distribution. This is then used to find the maximum a posterior probability (MAP) estimate of the signal. In calculating the a posteriori joint distribution further important qualities can be derived including the marginal posterior distribution of each individual change-point and parameter, along with useful information like the probability that there are $x$ number of change-points in the observed data.

The observed data from the experiments are the unobserved signal of increased intensities of light in the experiment due to the proton pump being active plus noise of random variations due to various different sources (e.g. camera noise, biological noise). Before initiation of activity by addition of ATP, single vesicle data contains a sample from which the variance of the noise and the base line around which the samples (on average) fluctuate is calculated. Without loss of generality, we consider the standardized case where the variance equals 1 and the baseline is at 0.

As the entire trajectory of the observed process in continuous time is not available, we operate with observations that are sampled at times $t = 1, 2, \dots, n$ (again for notational simplicity). Thus the observed data are $\boldsymbol{X} = (X_1, X_2, \dots, X_n)$ . As there is a finite set of observed data there is some probability only the first few (or indeed none) of the change-points considered are in this set. Hence the method is robust so long as there are less change-points present in the data than the number of change-points allowed in the model.

The likelihood function $L$ defined below can be seen as an approximation of the continuous time likelihood function using the discrete observed data

$$L(\boldsymbol{\theta}, \boldsymbol{\nu} | \boldsymbol{X}) = \prod_{t=1}^{n} \exp(\mu(t, \boldsymbol{\theta}, \boldsymbol{\nu}) X_t + 0.5\, \mu(t, \boldsymbol{\theta}, \boldsymbol{\nu})^2) .$$

Equivalently, this can also be derived by using a discrete time model for the signal. Given the joint probability density function $f(s)$ of the mode-switching events $\boldsymbol{\theta}$ and the joint probability density function $g(v)$ of the parameters $\boldsymbol{\nu}$ (see the previous section), the unnormalized a posteriori joint distribution for estimating $\boldsymbol{\theta}, \boldsymbol{\nu}$ can be calculated as follows $L(s, v | \boldsymbol{X}) f(s) g(v)$. Then the MAP estimate $\widehat{\boldsymbol{\theta}}, \hat{\boldsymbol{\nu}}$ of the mode-switching events and parameters are given by the values that maximize the unnormalized a posteriori joint distribution. Finally, these estimates give the MAP estimate of the signal to be $\mu(\mathrm{t}, \widehat{\boldsymbol{\theta}}, \hat{\boldsymbol{\nu}})$.

Using the normalized a posteriori distribution and integrating across all but one variable gives its marginal distributions (for example the a posteriori distribution of the first change-point). Given observed data $\boldsymbol{X}$, we can then find the probability that the first mode-switch event is present in this dataset.

Ideally the range of the distributions for $\alpha$, $\beta$ and $\gamma$ would be as large as possible. However, due to numerical constraints, for the application of the method of Bayesian stochastic filtering on the real data we used the following constants to calculate the MAP estimates: The parameter of the exponential distributions of the mode-switching events is $10^{(-5)}$, $\alpha$ is in the range $\max(\boldsymbol{X}) - 2$ to $\max(\boldsymbol{X}) + 4$, and $\beta$'s and $\gamma$'s are in the range 0.01 to 3.93.

**Sliding method**

It can be seen that increasing the possible number of change-points in the problem increases the dimension of the joint distribution and so the numerical complexity of producing the MAP estimates. It is not known a priori how many mode-switch events will be present in a sample of observed data, so one option would be just to choose a number of mode-switch events that is large enough that the Bayesian stochastic filtering returns an estimate in which not all of the change-points looked for are observed in the data. However, even for a relatively small number of change-points (≥ 5) the numerical calculations can become highly time consuming, even with very efficient C++ code.

As MAP estimates were needed to be produced for a large number of samples, a novel sliding method of Bayesian stochastic filtering was developed which aimed to first segment the data into regions where there is only one mode-switch event and then produce the MAP estimate for these segments individually. This also involved updating our distribution of the vesicle equilibrium level $\alpha$. This greatly reduced the complexity of required in the Bayesian stochastic filtering and made the method tractable for these large sets of data (see Extended Data Table 1 for a description of the code).

**Calculation of proton-pumping mode probabilities**

The probability that the V-ATPase is found in a proton-pumping mode was calculated using $P_{pumping} = \sum t_{on} / (\sum t_{on} + \sum t_{off})$, except for Fig. 4d, where the probability as a function of ATP was estimated as $P_{pumping} = \sum t_{on} / t_{ATP}$, where $t_{ATP}$ is the total time of the recording post ATP injection.

**Calculation of the transition energy barrier heights for on-to-off and off-to-on cycle transitions.**

Transition rates (main Fig. 3c, Extended Data Fig. 11) were calculated using transition state theory, where the pumping and inactive modes are considered ground states separated by an activation barrier (the transition state). According to transition state theory the energy required to achieve the transition state (and cross the barrier) is:

$$\Delta G^{\ddagger} = -RT\ln\frac{hk_{i,j}}{k_B T}$$

where R is the gas constant, T is the absolute temperature, $k_B$ is a Boltzmann's constant, h is Planck's constant, $k_{i,j}$ is the measured transition rate from pumping to inactivation or from inactivation to pumping states. Under the assumption that we have a two-mode model, proton-pumping and inactive mode rate constants can be calculated using equations

$$k_{on,off} = \frac{1}{\langle t_{on} \rangle}$$

$$k_{off,on} = \frac{1}{\langle t_{off} \rangle}$$

where $\langle t_{on} \rangle$ and $\langle t_{off} \rangle$ are the average dwell times in pumping and inactive modes for the V-ATPase.

**Calculation of free energy difference between proton-pumping and inactive modes.**

We calculated Gibbs free energy difference between proton pumping and inactive modes for different electrochemical conditions (Extended Data Fig. 11) using equation,

$$\Delta G = -RTln \frac{P_{pumping}}{1-P_{pumping}}$$

where $P_{pumping}$ and 1-$P_{pumping}$ represent the probabilities of the pump being in proton-pumping or inactive modes respectively.

**Effects of membrane potential on mode-switching**

The regulatory effects of $\Delta\Psi$ on $\tau_{on}$ and $\tau_{off}$ cancelled out each other, so that the overall probability of being in a proton-pumping mode remained independent of $\Delta\Psi$ within the experimental uncertainties, ($P_{on}^{+Cl}$ = 0.59 ± 0.11 and $P_{on}^{-Cl}$= 0.54 ± 16, Extended Data Fig. 11b). Notably, because the electrochemical gradient collapses back to zero during the periods of inactivity it cannot affect directly $\tau_{off}$, thus the regulation of $\tau_{off}$ is likely the result of a molecular memory effect mediated via the proton pumping mode.

**Concentration of active V-ATPases during microscopy measurements**

After immobilization of LUVs and fusion with SVs, there are ~100 $SV_h$ in a 0.0064 $mm^2$ FOV. Given that the area of the microfluidic chamber is 64.6 $mm^2$ and that there are 1.2 V-ATPases/$SV_h$, the total number of V-ATPases in the chamber is $1.2 \cdot 10^6$. Hence, given the chamber volume, 30 µl, the concentration of V-ATPases present during the course of an experiment is ~70 fM.

#### Supplementary Methods

**Image analysis**

Data acquired from light microscopy experiments were analyzed using a custom software suite developed in the lab as well as custom scripts in MATLAB. A single particle localization algorithm identified single vesicles and their corresponding locations in the FOV. A drift correction algorithm was implemented whenever thermal fluctuations caused XY drifts during recordings[2]. Once image sequences have been stabilized and particles were identified[3], we defined 5x5 pixel ROIs around each particle and got the integrated intensity readout by summation of intensity values of individual pixels. This then applied throughout the image time series and therefore, single vesicle kinetics were produced. After each activity measurement, a stack of 20 images at saturating pH conditions was acquired to assist in the conversion of intensity to pH via calibration curves (for more details see section: Calibration of pH in vesicles). For pHrodo-PE the signal saturated at approximately pH 3 (Extended Data Fig. 7).

Normalized intensity is defined as the raw intensity extracted divided by the mean intensity of the baseline, $I_{norm} = I_{raw}/I_{base}$. Standardized intensity is defined as the raw intensity reduced by the mean baseline intensity and divided by the standard deviation of the signal at the base line ($I_{std} = (I_{raw}-I_{base})/\sigma_{base}$). Normalized intensities were used as a typical representation of signals throughout this article. Standardized intensities were used by the stochastic filtering algorithm.

Active transport kinetics were initially identified via threshold filtering based on the signal-to-noise (SNR) of each kinetic trace. *SNR* is defined on an individual vesicle basis as the maximum normalized intensity reduced by one and divided by the normalized standard deviation of the baseline, $SNR = (I_{norm,max}-1)/\sigma_{norm,base}$. The selection of active transport kinetics was then manually classified based on the number of steps (intensity plateaus) identified (Fig. 1d, e). Only data with a single intensity step, corresponding to single transporter activity, was considered for single molecule data presented in the present article. Single vesicle activity traces that showed no stochastic behavior were not used in the single molecule data.

**Dwell time analysis**

A stochastic filtering algorithm (see Supplementary Discussion) was used to identify mode-switching events in single molecule data. In cases where there was overdetection of events, postprocess filtering algorithms were applied to clean up event data. Less than 10% of event data was subjected to postprocessing. A minor fraction of event detection data (less than 5%) was manually fine-tuned.

In order to calculate dwell times, we only considered single molecule data that showed at least one stochastic event. Proton-pumping dwell times ($t_{on}$) were calculated as the duration from the onset of acidification up until the point when the signal starts decreasing again which corresponds to the V-ATPase switching off (Fig. 2b). Inactive dwell times ($t_{off}$) are calculated as the duration from the start of signal decrease until the next on-cycle event (Fig. 2b). Dwell times of modes were only included in this analysis from the onset of the first proton-pumping event and were not interrupted by the ending of the recording. The probability that the V-ATPase is found in a pumping mode was calculated as the total time the V-ATPase spent in a pumping dwell versus the total time spent in both in pumping and inactive dwells post ATP injection ($P_{pumping} = \Sigma t_{on}/\Sigma(t_{on} + t_{off})$). Any event prior to the first pumping-dwell or interrupted by the end of the recording is rejected as the true duration of those events is unknown.

**Calibration of pH in vesicles**

In order to calibrate activity traces, a semi-local calibration methodology was followed (See also Extended Data Fig. 7). Initially, we used DOPE-pHrodo labelled liposomes with the same lipid composition as mentioned previously (DOPC:DOPS:Chol:(18:1 Biotinyl Cap PE):DOPE-pHrodo 64.4:10:25:0.5:0.1) rehydrated in calibration buffer (150 mM K-Gluconate, 2 mM MOPS and 2 mM $MgSO_4$) and adjusted with Tris to reach a pH of 7.23. These liposomes were immobilized according to protocols mentioned above and they were incubated in calibration buffer which additionally included 60 nM valinomycin (val) and 5μM Carbonyl cyanide m-chlorophenyl hydrazone (CCCP), to facilitate unconstrained influx of protons and efflux of potassium ions. Subsequently they were treated with calibration buffer at nine different pH values ranging from 7.23 to 3.19. For the calibration buffers, pH was adjusted with citric acid. For each pH measured, a stack of 20 images was recorded. Four replicates of this experimental process were performed.

Using the custom software suite, we identified vesicles and extracted raw intensities for the different pH values recorded. For each vesicle the calibration data was fitted with the sigmoidal function:

$$I(pH) = I_{base} - \frac{I_{max}}{1+e^{(pKa-pH)/rate}} \qquad (1)$$

$I$ is the raw intensity recorded, $I_{base}$ is the lower asymptote, $I_{max}$ is the upper asymptote, $pK_a$ is the acid dissociation constant and the *rate* corresponds to the sharpness of the sigmoidal curve. Each fitted sigmoid then was normalized to the intensity corresponding to pH 7.1 as that corresponds to the pH of the baseline intensity of activity traces. We, then, extracted the mean values of the distributions of pKa and rates. As the standard deviations of these parameter distributions are narrow the vesicle population displayed a well-defined and homogeneous behavior for these parameters. $I_{base}$ and $I_{max}$, which define the dynamic range of the fluorophore, are more heterogeneous and thus are locally defined for each single $SV_h$ after its activity has been acquired.

After recording single vesicle acidification kinetics in the activity buffer at pH 7.1, we incubate the $SV_h$ for five minutes in the activity buffer, but at a signal saturating pH 2.85 in the presence of 5 µM CCCP and 30 mM choline chloride. Chloride is added to negate any membrane potential being established by proton influx. The absence of any other species of counterions (or neurotransmitters), such as potassium or sodium, ensures that secondary transporters, found on the vesicle membrane, cannot transport substrates that could alter measuring conditions. Then, a sequence of twenty images is acquired under these conditions.

Calibration of activity data is done by first normalizing the raw traces. This is done by subtracting half of the baseline signal from the whole time series to correct for the fact that any signal variation reported due to activity of the V-ATPase is stemming from the pHrodo-PE lipids of the inner leaflet of the bilayer. Then, the trace is normalized to the new baseline intensity. Additionally, the raw intensity, $I_{sat}$, at saturating conditions (pH 2.85) is divided by two to compensate for contribution of pHrodo from both leaflets and then normalized to the baseline intensity. Solving equation (1) for pH we end with an expression of pH as a function of intensity and $I_{base}$ and $I_{max}$ are determined from the conditions pH = 7.1 when $I = 1$ and pH = 2.85 when $I = I_{sat}$. It is assumed throughout this analysis that pHrodo-PE lipids are equally distributed between the two leaflets of the bilayer.

**Proton leakage in hybrid vesicles and LUVs**

We performed separate proton leakage experiments and also monitored leakage lifetimes during activity measurements in order to determine and deconvolve the leakage mechanisms. To determine passive leakage through the membrane, we immobilized LUVs on the glass surface and subsequently added SVs to initiate in situ fusion as described earlier. Subsequently, $SV_h$ and LUVs are incubated for approximately 10 minutes in activity buffer (300 mM glycine, 2 mM MOPS and 2 mM $MgSO_4$) at pH 6.2 (pH is adjusted with citric acid). Then we injected activity buffer at pH

7.1. We record pH jumps from pH 6.2 to 7.1 and monitor the leakage kinetics by means of fluorescence signal reduction (Extended Data Fig. 4). In order to distinguish between LUVs and $SV_h$, upon completion of the leakage measurement, we added 1 mM ATP and monitored activity thus allowing us to distinguish the two vesicle populations.

Once leakage data was collected for both LUVs and $SV_h$ we fitted the single vesicle kinetics from the moment we did the buffer exchange with a double exponential function. The outer leaflet will respond fast to the buffer exchange, subject only to the flow rate induced by the peristaltic pump used to exchange media. Thus, the fast component of the fit corresponds to fluorescence stemming from the outer monolayer. Passive leakage through the vesicle membrane will be reported by the slow component of the double exponential. Leakage lifetimes of $SV_h$ and LUVs were found to be in excellent agreement between them, strongly suggesting that no other protein on the membrane of $SV_h$ provides a pathway for protons to leak under our experimental conditions. Moreover, the fact that no stochastic jumps were observed during the pH jumps for either vesicle population indicates that any observed off-cycle modes are solely attributed to the V-ATPase. It is also noteworthy that no transprotein leak was observed for the V-ATPase in the absence of catalytic substrate or upon inhibition with bafilomycin.

Leakage lifetimes of $SV_h$ during activity (in the presence of ATP) were estimated by fitting leakage rates with a mono exponential function when the V-ATPase switched to an inactive or leaky mode. Slow decay is attributed to membrane leakage while the fast decay stems from a transprotein leakage pathway through the V-ATPase (Fig. 2e). Due to the transprotein leak being 10 times faster than passive leakage through the membrane, a mono exponential function is sufficient to describe fast efflux kinetics. Lastly, single molecule leakage rates upon addition of 200 nm of bafilomycin were fit in a similar manner (Fig. 2e).

#### References

1. Bain, A. & Crisan, D. *Fundamentals of stochastic filtering*. (Springer, 2009).
2. Guizar, M. Efficient subpixel image registration by cross-correlation (https://www.mathworks.com/matlabcentral/fileexchange/18401-efficient-subpixel-image-registration-by-cross-correlation), MATLAB Central File Exchange. Retrieved September 28, 2021. (2021).
3. Furst, E. M. Particle tracking with Matlab. (2015).
