## Extended Data Figures for "Pump-Rest-Leak-Repeat: regulation of the mammalian-brain V-ATPase via ultra-slow mode-switching"

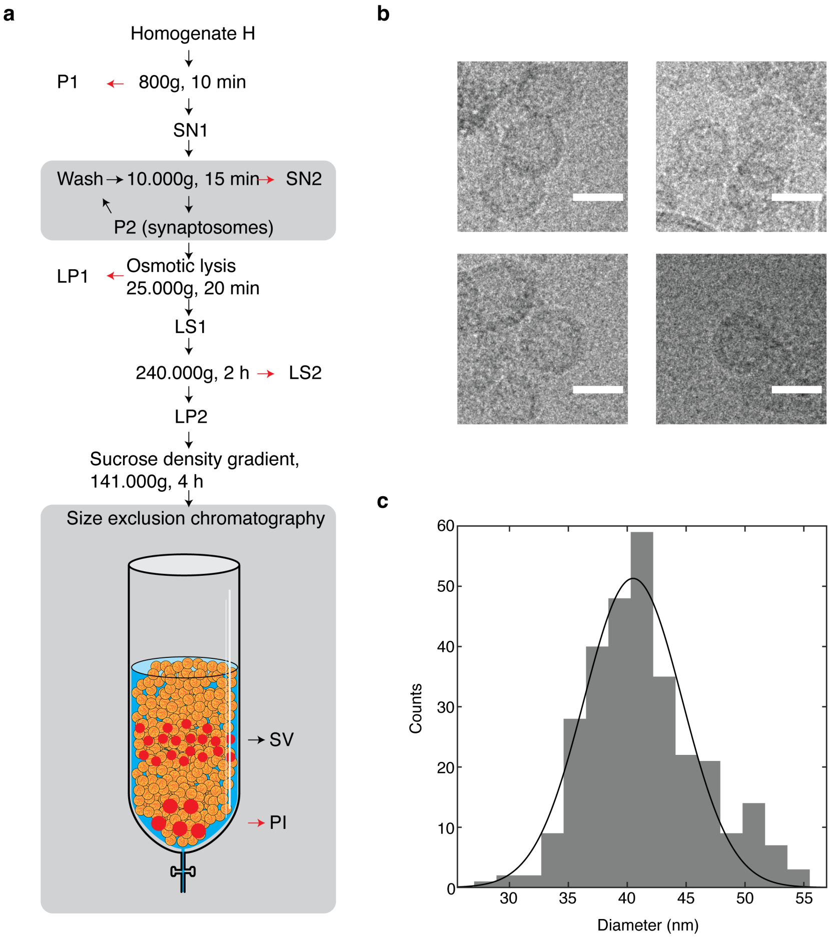


**Extended Data Fig. 1 | Purification and morphology of SVs.**

**a**, Protocol diagram for the purification of SVs as adapted from a previous publication[1]. **b**, Cryo-electron micrographs of purified synaptic vesicles. SV fraction was highly pure with larger organelles rarely found in the micrographs. Scale bar is 40 nm. **c**, Distribution of SV diameters from cryo-electron micrographs. Population of SVs was found to be monodisperse with a highly conserved vesicle size.

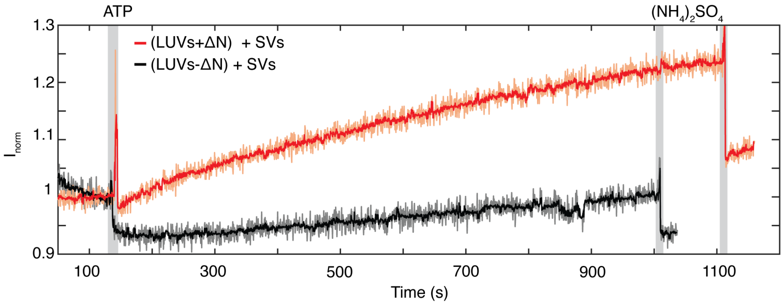


**Extended Data Fig. 2 | Macroscopic ensemble experiment of activity**.

Proton pumping was initiated by addition of 2 mM ATP and stopped upon addition of 20 mM ammonium sulfate. Proteoliposomes reconstituted with the ΔN complex were able to fuse with SVs and produce $SV_h$ that contain the V-ATPase (red trace). No fusion and therefore no activity was observed for liposomes lacking the ΔN acceptor complex (black trace). Raw data (orange and grey trace) was processed with a Chung-Kennedy filter (red and black trace)[2]. All subsequent acidification traces shown in figures have been processed with a Chung-Kennedy filter, unless stated otherwise.

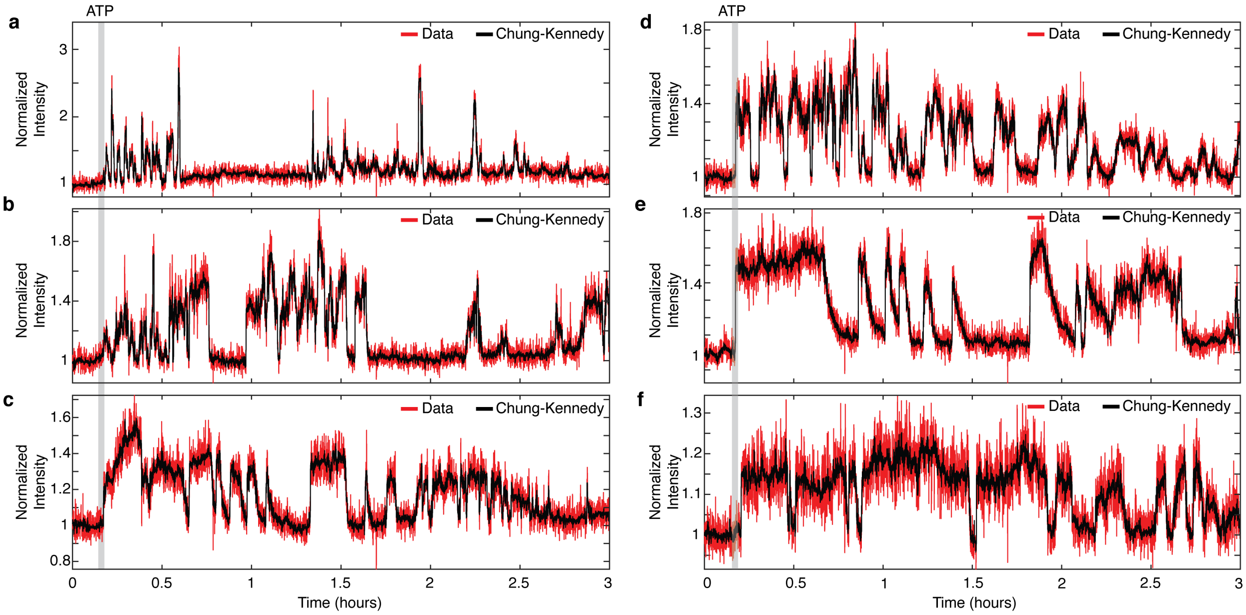


**Extended Data Fig. 3 | Ultra-long-term activity recordings of single V-ATPases in single $SV_h$.** Activity recordings that spanned the duration of three hours revealed mode-switching of single V-ATPases. **a-f**, Representative single molecule acidification kinetics displaying long-lived proton pumping and inactive modes. Activity was initiated upon addition of ATP. Red traces correspond to raw data (not corrected for photobleaching) that have been normalized to the baseline. Black traces correspond to data processed with a Chung-Kennedy filter.

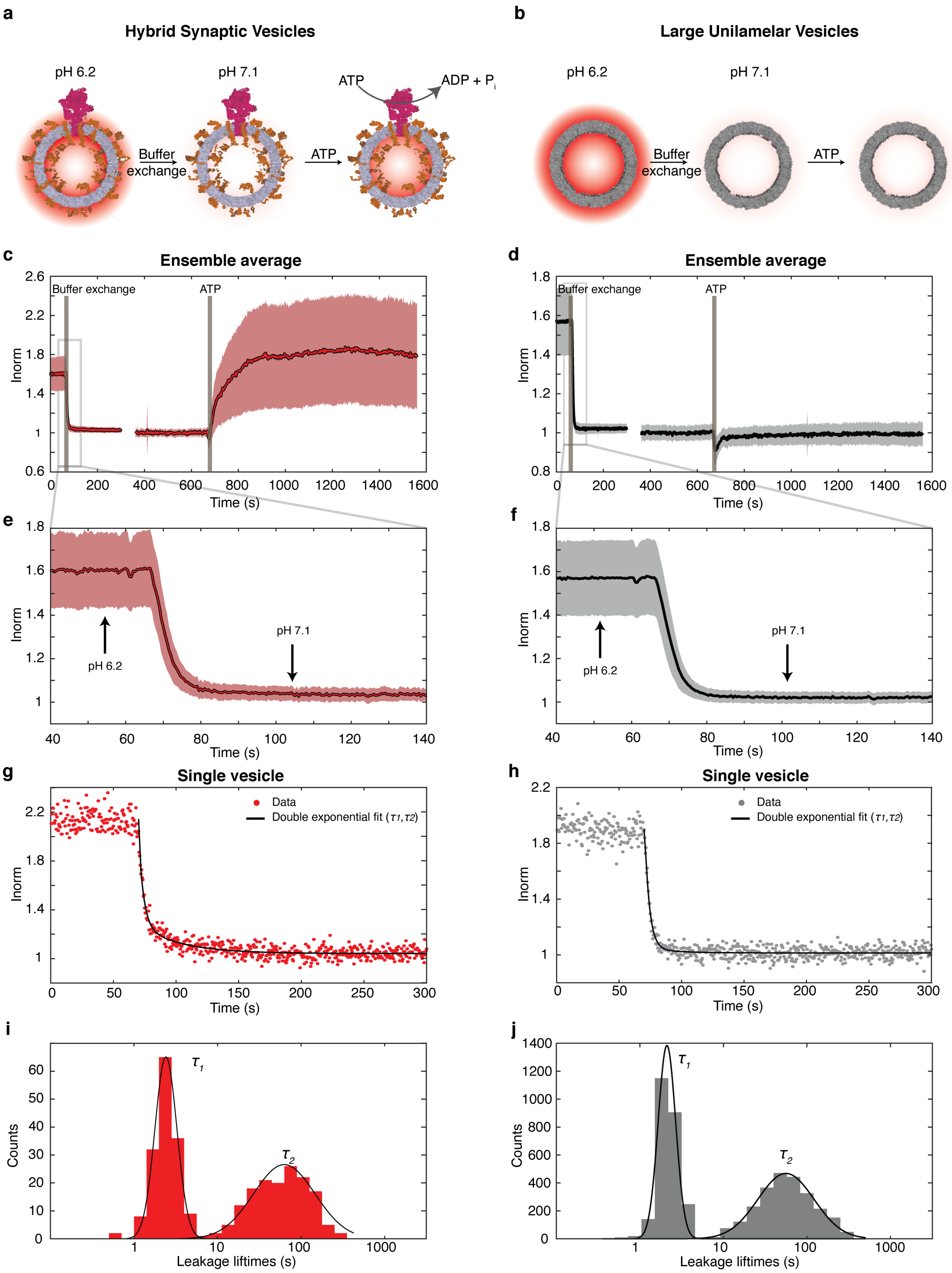


**Extended Data Fig. 4 | Experimental determination of passive proton leakage in $SV_h$ and LUVs.**

Left panels correspond to data from $SV_h$ while right panels correspond to data from LUVs. **a**, **b**, Schematic illustrations of the experimental procedure. $SV_h$ and LUVs are incubated in a glycine-

MOPS buffer at pH 6.2. Subsequently, we exchange the buffer with a glycine-MOPS buffer at *pH* 7.1 and record the change in signal intensity. Lastly, we inject ATP and record activity to differentiate between $SV_h$ containing the V-ATPase and LUVs that don't respond to ATP. During buffer exchange, pHrodo lipids in both leaflets are subjected to the different *pH*, while during injection of ATP, only the inner leaflet of $SV_h$ will report changes in *pH*. **c-f**, Ensemble average traces for *pH* jumps and injection of ATP. Lines correspond to the mean of the ensemble while shaded areas correspond to one standard deviation of the spread of the single vesicle data. **g**, **h**, Typical single vesicle kinetics of signal decrease. Data is fitted with a double exponential function to extract lifetimes of fluorescence decay for the outer leaflet (fast) and the inner leaflet (slow). **i**, **j**, Histograms of lifetimes derived from the double exponential fits on the single vesicle data. Gaussian fits on these histograms produced average fast and slow values. The fast component corresponding to the outer leaflet was found to be $\tau_1$ = 2.41 s ± 1.02 s and $\tau_1$ = 2.11s ± 1.02 s for $SV_h$ and LUVs respectively. The slow component corresponding to the inner leaflet and tracks proton leakage lifetimes was found to be $\tau_2$ = 62.95 s ± 1.07 s and $\tau_2$ = 56.10 s ± 1.03 s for $SV_h$ and LUVs respectively. Leakage lifetimes were smoothly monotonic over time indicating a single mechanism for proton efflux.

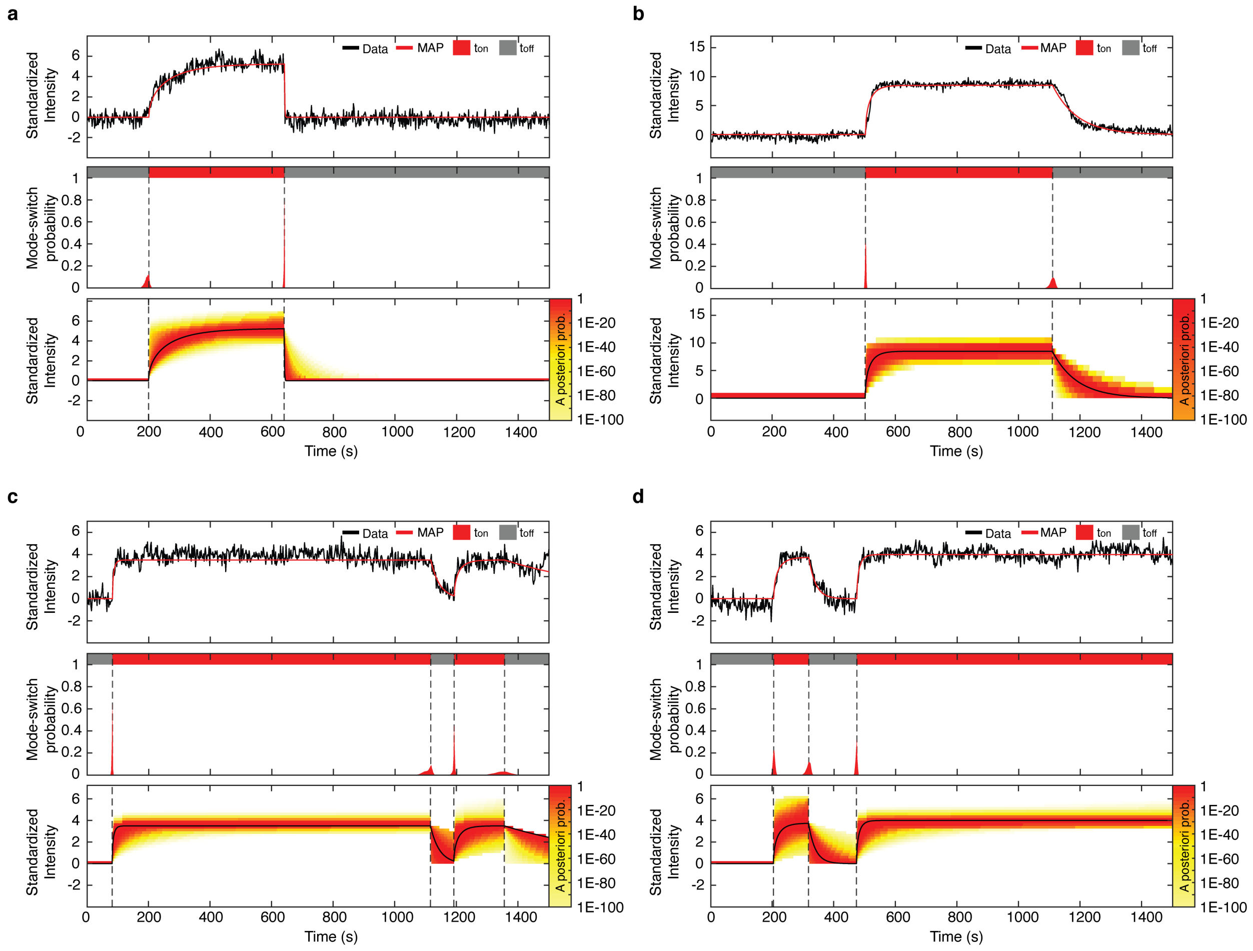


**Extended Data Fig. 5 | Detection of mode-switching events and MAP fits with an exponential model.**

**a-d**, Examples of single molecule data being analysed using stochastic filtering techniques and exponential model. The algorithm is run sequentially along observed data and then segments the data into areas where the proton pump is found in either proton-pumping or inactive states. These areas are determined using the posterior distributions for mode-switching events seen as the red distributions in the ‘mode-switching probability’ plots. When there is enough confidence (99% certainty) that an event has been observed, the data is segmented at the event time with the highest probability (in terms of the joint distribution, see Stochastic filtering section in supplementary text), shown with dashed lines. Stochastic filtering techniques are then used on the segmented data to find the parameters with the highest likelihood given the exponential model. The heatmap represents the (a posteriori) probability of where the signal is deemed to be, given the observed data and the exponential model. The MAP estimate of the signal is seen in these heatmap plots in black.

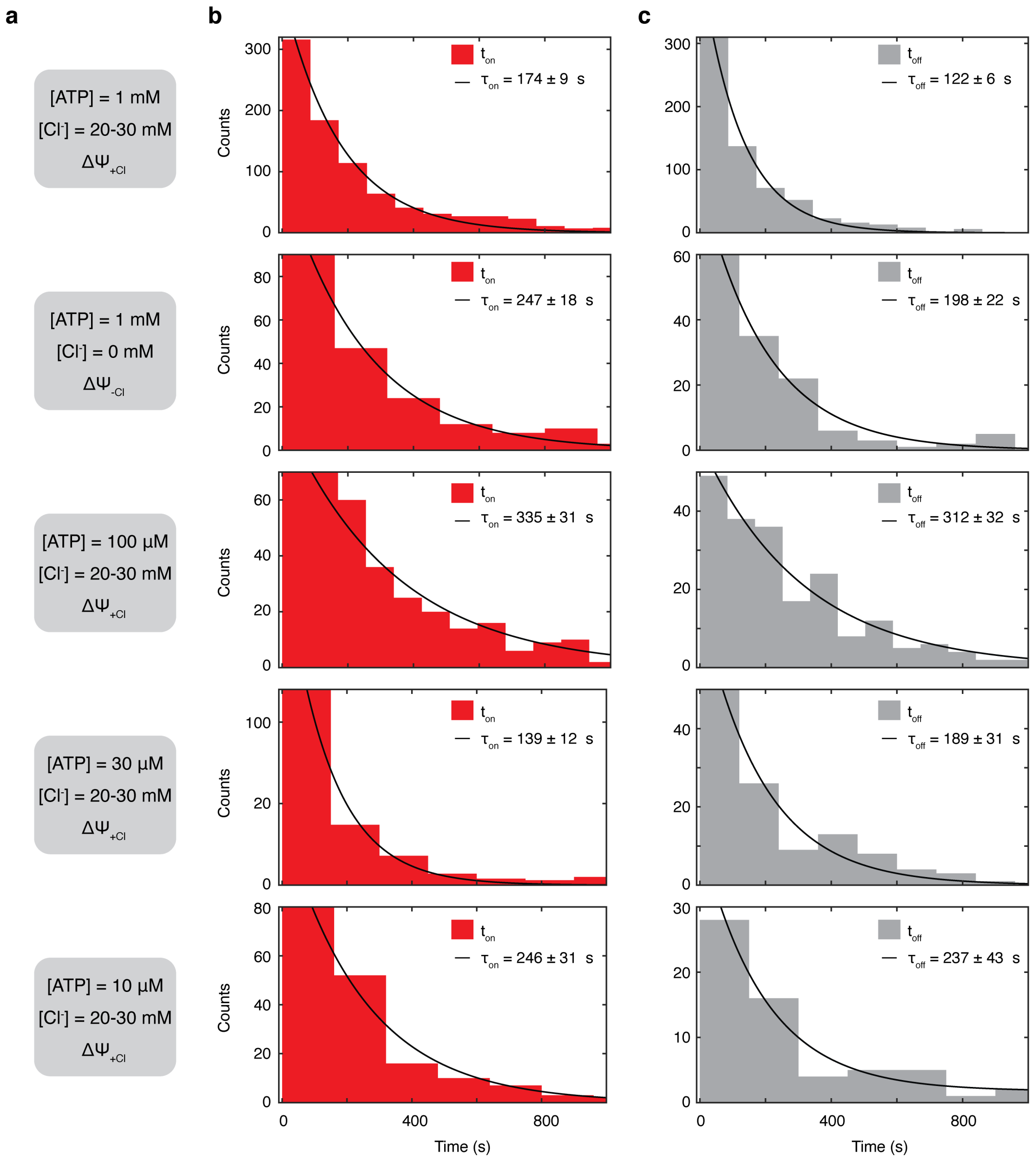


**Extended Data Fig. 6 | Dwell times of the proton-pumping and inactive modes for the V-ATPase in the presence/absence of chloride and different concentrations of ATP.**

Each line corresponds to data collected from different sets of experiments. **a**, Experimental conditions during which dwell times of states of the V-ATPase were extracted. **b,** Distributions of proton-pumping dwell times. **c,** Distributions of inactive dwell times. Note: any events that were interrupted by the end of the recording, were discarded. Only inactive modes after the initiation of the first pumping event were considered in the data. *n* = 4-8.

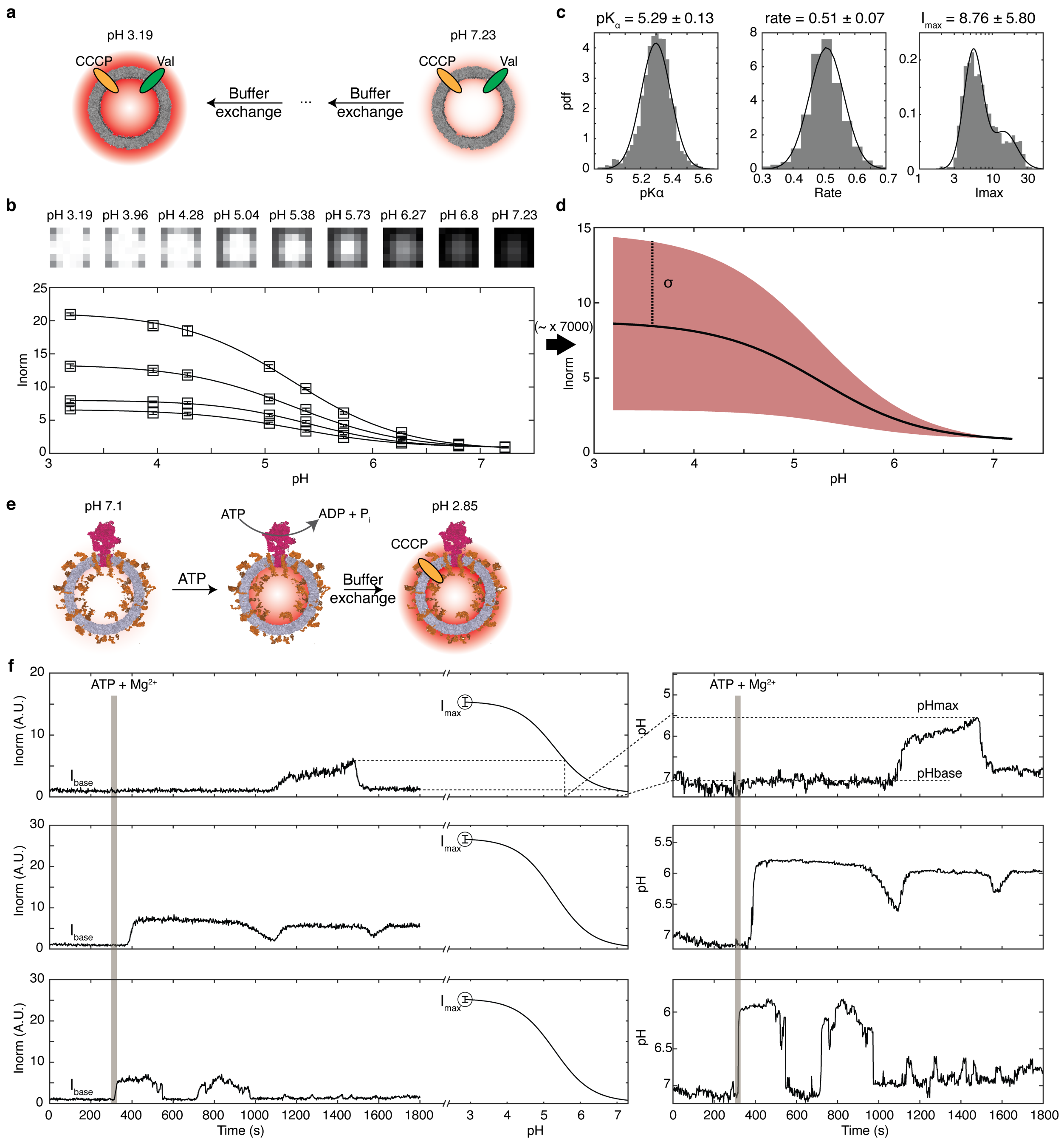


**Extended Data Fig. 7 | pH calibration methodology for single molecule activity data.**

**a**, Schematic illustration of pH calibration experiments. LUVs are incubated in a K-gluconate-MOPS buffer at *pH* 7.23 in the presence of valinomycin and CCCP to equilibrate chemical gradients. Subsequent injection of the same buffer (including valinomycin and CCCP) at decreasing *pH* concentrations leads to equilibrated acidification of both the lumen and the extraluminal space around the vesicles. The *pH* changes are reported as an increase in fluorescence. **b**, Representative examples of single vesicle calibration data. Data has been normalized to the activity baseline value (see supplementary text). Each vesicle reports a change in fluorescence as a function of *pH* which is then fitted with a sigmoidal function. Error bars represent the SD of fluorescence intensity for a stack of 20 images. **c**, Distributions of the sigmoidal fit parameters. The dissociation constant and growth rate of the vesicles show a normally distributed population with very narrow standard deviations. The fluorescence, $I_{max}$, at saturating *pH*, however showed a larger spread. Thus, the dissociation constant and the growth rate are globally determined from these experiments while the $I_{max}$ saturation value is locally determined for $SV_h$ post activity. **d**, Ensemble average of sigmoidal fits for the entire population of vesicles. Black line corresponds to the mean while red-shaded area corresponds to the standard deviation of the population. **e**, Illustration of the typical process during activity measurements. After recording a baseline, ATP is injected into the system and V-ATPase activity is triggered. After the end of the activity recording, the $SV_h$ are treated with activity buffer at signal saturating *pH* 2.85 including CCCP and chloride to allow for influx of protons and chloride counterions to determine Imax locally for each vesicle. **f**, Example activity traces and their corresponding calibration curve determined by the global dissociation constant, the global growth rate (determined in panel (**c**)), the local baseline and saturation values, $I_{base}$ and $I_{max}$ respectively. Henceforth, normalized intensities of activity measurements can be mapped onto corresponding *pH* values, as shown in the right-hand side traces. Error bars represent the SD of fluorescence intensity for a stack of 20 images.

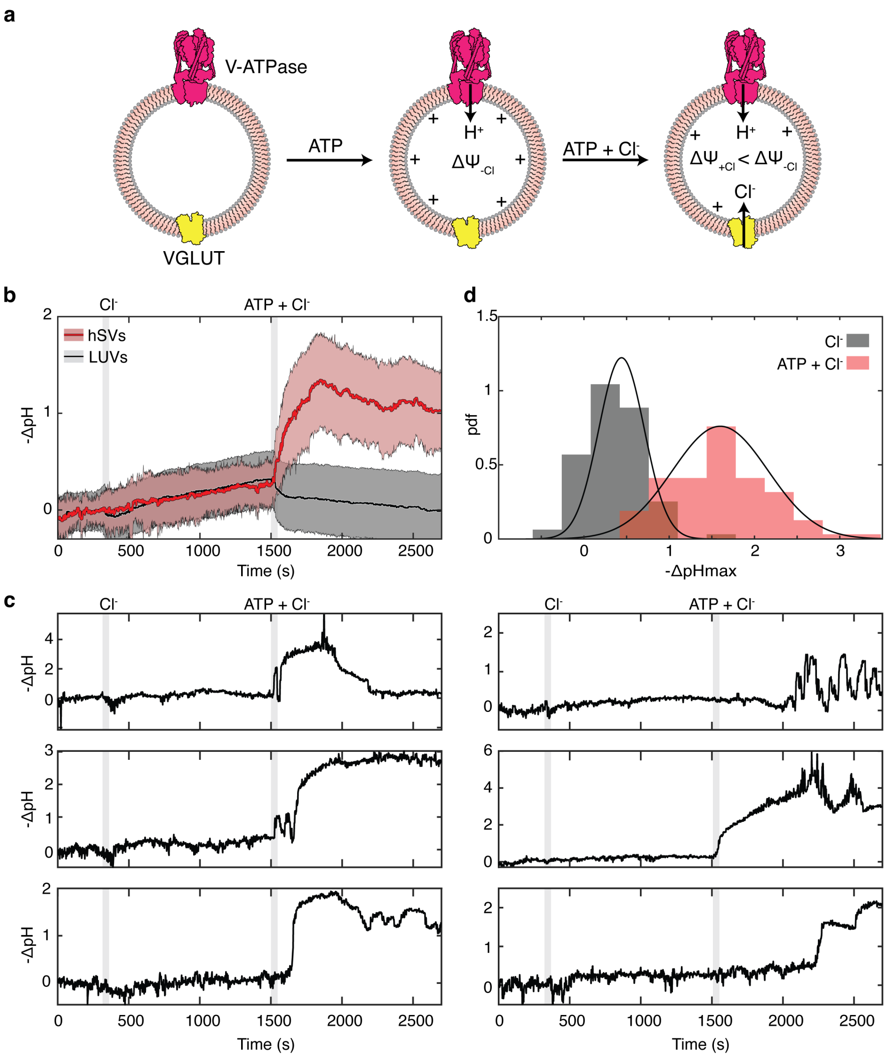


**Extended Data Fig. 8 | Luminal chloride reduces membrane potential and induces mild acidification in $SV_h$ without introducing stochastic effects.**

**a**, Schematic illustration of membrane potential buildup during activity measurements. Upon addition of ATP, protons are pumped into the vesicles and an inside positive $\Delta\Psi_{-Cl}$ is established. When extraluminal chloride is introduced, $Cl^-$ is transported into the lumen by VGluT and partially releases the potential resulting in $\Delta\Psi_{+Cl} < \Delta\Psi_{-Cl}$. **b**, Effects of chloride on ensemble average acidification kinetics of $SV_h$ and LUVs. After recording the baseline, 30 mM choline chloride are introduced into the system. Subsequently, 30 mM choline chloride and 1 mM $Mg^{2+}$ and ATP are added, and for $SV_h$, V-ATPase specific activity is monitored. Minor acidification was observed upon addition of chloride for both LUVs and $SV_h$, which is attributed to passive permeation of chloride ions through the membrane. No acidification was observed for LUVs upon addition of ATP. **c**, Typical single molecule traces for $SV_h$ showcasing the behavior of the pump under these conditions. Addition of chloride only does not introduce any discernible stochastic effects in the form of acidification spikes. Contrary, addition of ATP introduces stochastic fluctuations in acidification levels. **d**, Histogram of the maximal acidification of $SV_h$, $-\Delta pH_{max}$, under the aforementioned conditions. When only chloride is present, the average $-\Delta pH_{max}$ = 0.35, while in the presence of both chloride and ATP, $-\Delta pH_{max}$ = 1.76. Black dots correspond to independent experiments performed.

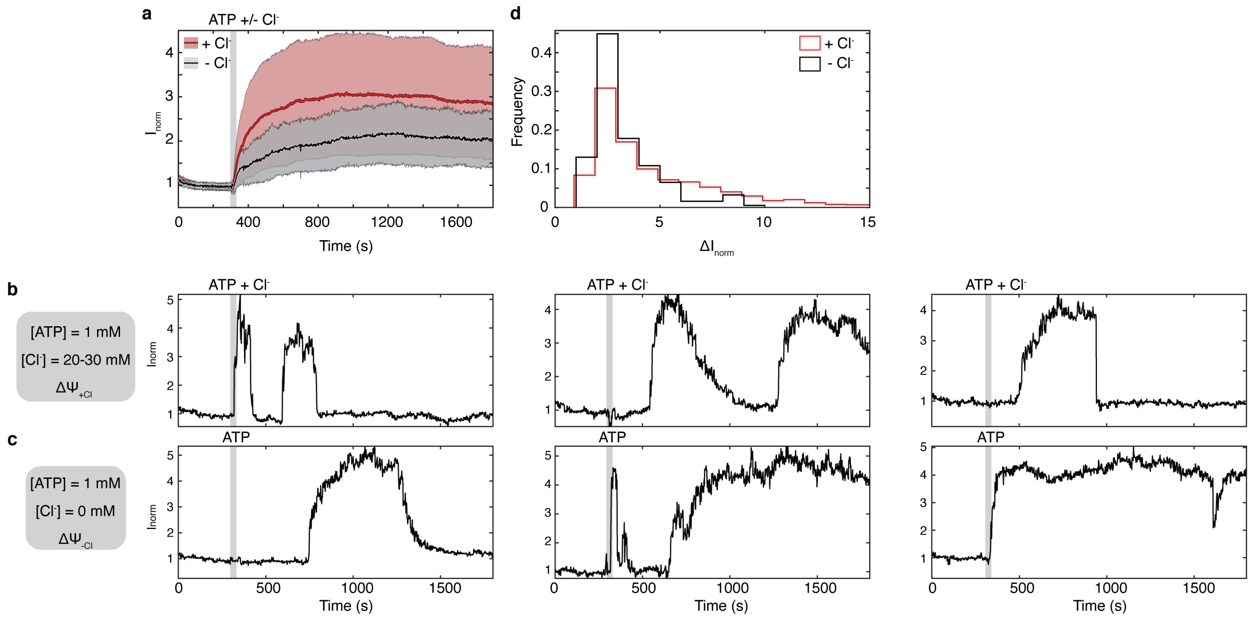


**Extended Data Fig. 9 | Single vesicle acidification kinetics in the presence and absence of chloride.**

Data analyzed in these experiments was used to determine dwell times for both proton-pumping and inactive modes and their relation to electrochemical gradients (Fig. 2c, d, Fig. 3d-f). **a**, Ensemble average acidification kinetics of $SV_h$. In the absence of chloride, a membrane potential clamps the activity of the V-ATPase (black line). When 30 mM choline chloride is introduced, the potential is released and the V-ATPase can establish larger gradients (red line) (also see Fig. 3a, b). **b**, **c**, Typical single molecule traces for the different conditions investigated in these experiments. **d**, Histograms of maximum intensities of single molecule data. Data shows different distribution of signals. The number of independent experiments were n = 8 for data in the presence of chloride (Panel (**a**), (**d**): Red data and Panel (**b**)) and *n* = 5 for data in the absence of chloride (Panel (**a**), (**d**): Black data and Panel (**b**)).

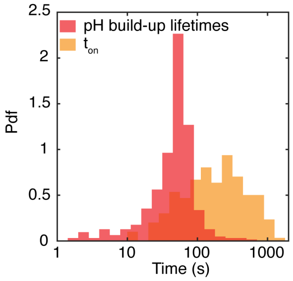


**Extended Data Fig. 10 | Comparison between kinetics of pH build-up and proton-pumping dwell times.**

Populations of *pH* build-up lifetimes and proton-pumping mode dwell durations. *pH* build-up data (red) was produced by fitting single exponentials on kinetic traces during periods of rise of activity. Data for $t_{on}$ (yellow) is taken from Extended Data Fig. 6b, second row and shown in a logarithmic scale.

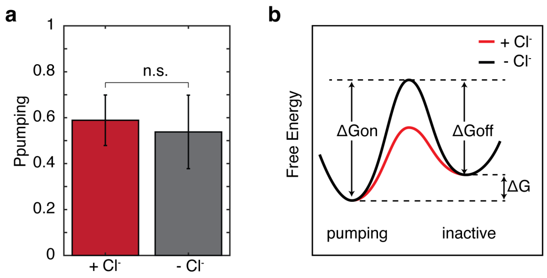


**Extended Data Fig. 11 | Electrochemical regulation of mode-switching dynamics.**

**a**, Proton-pumping mode probabilities in the presence or absence of chloride. No statistical significance was detected between the two populations. Error bars correspond to standard deviation of the mean between experiments. Total number of independent experiments is n = 8 and n = 5 for $\Delta\Psi_{+Cl}$ and $\Delta\Psi_{-Cl}$ respectively. **b**, Representation of the free energy landscape for the two different electrochemical conditions. The forward activation barriers, $\Delta G_{on}^{+Cl} = 86.2\ \mathrm{kJ \cdot mol^{-1}} \pm 0.3\ \mathrm{kJ \cdot mol^{-1}}$ and $\Delta G_{on}^{-Cl} = 86.7\ \mathrm{kJ \cdot mol^{-1}} \pm 0.4\ \mathrm{kJ \cdot mol^{-1}}$, are higher than those for backwards activation, $\Delta G_{off}^{+Cl} = 85.3\ \mathrm{kJ \cdot mol^{-1} kJ \cdot mol^{-1}} \pm 0.6\ \mathrm{kJ \cdot mol^{-1}}$ and $\Delta G_{off}^{-Cl} = 86.3\ \mathrm{kJ \cdot mol^{-1}} \pm 1.3\ \mathrm{kJ \cdot mol^{-1}}$, highlighting the fact that the probability the V-ATPase is found in a proton-pumping mode is higher. Additionally, both forward and backwards activation barriers were lower when the membrane potential was released which indicates that the frequency of transitions between modes is higher (see also Extended Data Fig. 6).

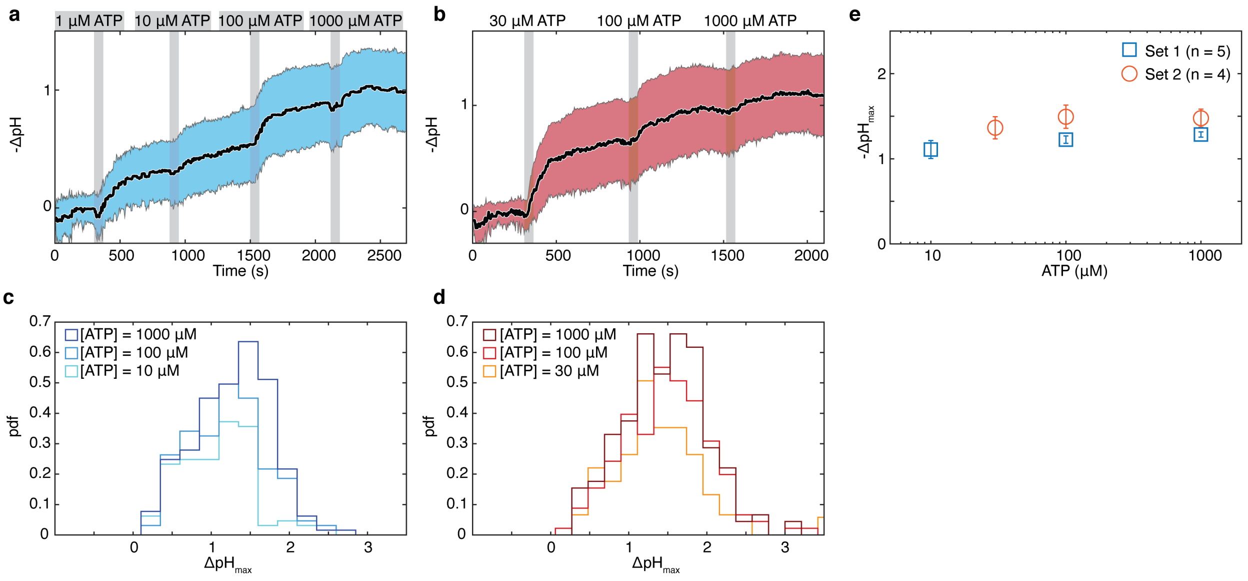


**Extended Data Fig. 12 | Additional data for ATP titration experiments.**

**a**, **b**, Ensemble average acidification kinetics for two different sets of experiments with varying ATP concentrations. Black lines correspond to the mean, while blue and red shaded areas correspond to standard deviation of the single vesicle data. Note: (**b**) is also shown in Fig. 4a. **c**, **d**, Population histograms displaying maximal acidification plateaus ($\Delta pH_{max}$) of the same single vesicles under different concentrations of ATP. In (**c**), maximum acidification of vesicles when [ATP] = 1 μM were not included. Pairwise two-sample Kolmogorov-Smirnov test showed for (**c**) statistical significance between [ATP] = 10 μM and either [ATP] = 100 (*) or [ATP] = 1000 (***). No statistical significance was observed between [ATP] = 100 μM and [ATP] = 1000 μM (n.s.). For (**d**), pairwise two-sample Kolmogorov-Smirnov tests showed no statistical significance for either of the pairs of ATP concentrations (n.s). **e**, Combined mean population values of $-\Delta pH_{max}$ of the histograms shown in (**c**) and (**d**) as a function of ATP concentration. Error bars correspond to SD of the mean of the independent experiments. A one-way ANOVA test gave a *P-value* < 0.05 (*) for set 1 (blue) and a *P-value* > 0.05 (n.s.) for set 2 (red). Note: data in Fig. 4c are the averaged values of set 1 and set 2 data combined.

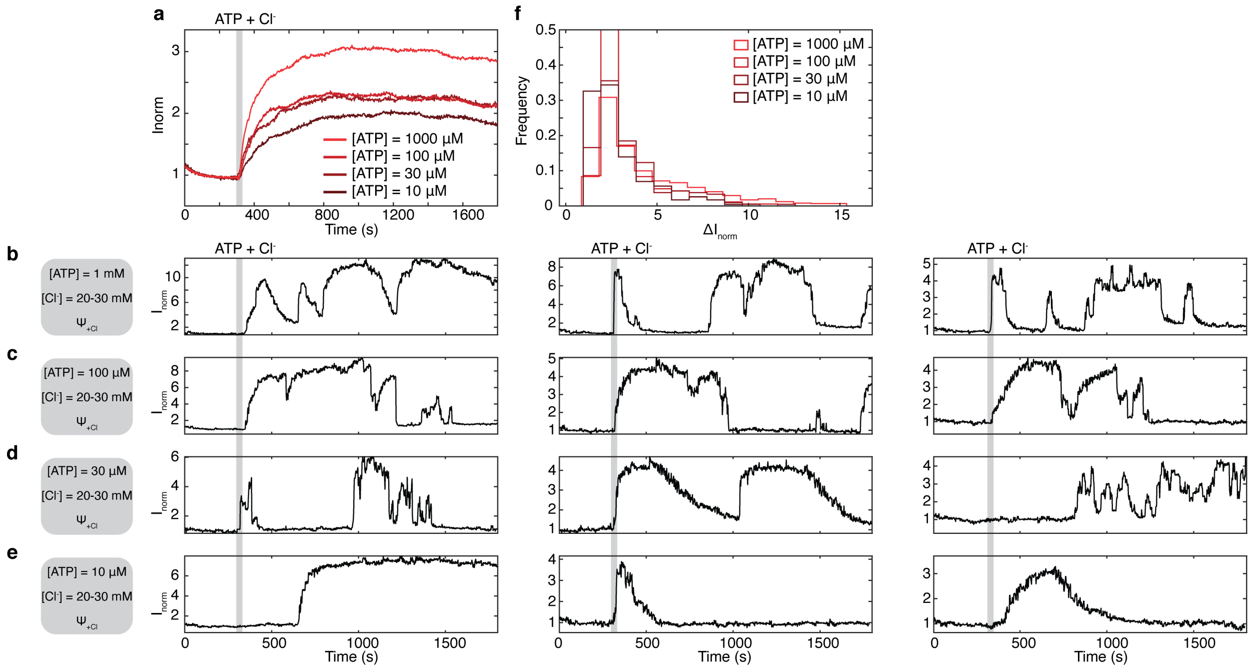


**Extended Data Fig | 13. Acidification kinetics under different catalytic substrate concentrations.**

Data analyzed from these experiments was used for calculating pumping-mode probabilities (Fig. 4d) and dwell times (Extended Data Fig. 6). **a**, Ensemble average acidification kinetics of $SV_h$ at different concentrations of ATP. Activity was initiated upon addition of ATP and 20-30 mM chloride. **b-e**, Typical single molecule traces at different concentrations of ATP (as stated in corresponding grey boxes). **f**, Histograms of maximum intensity of single molecule data. Total number of experiments was $n$ = 4-8.

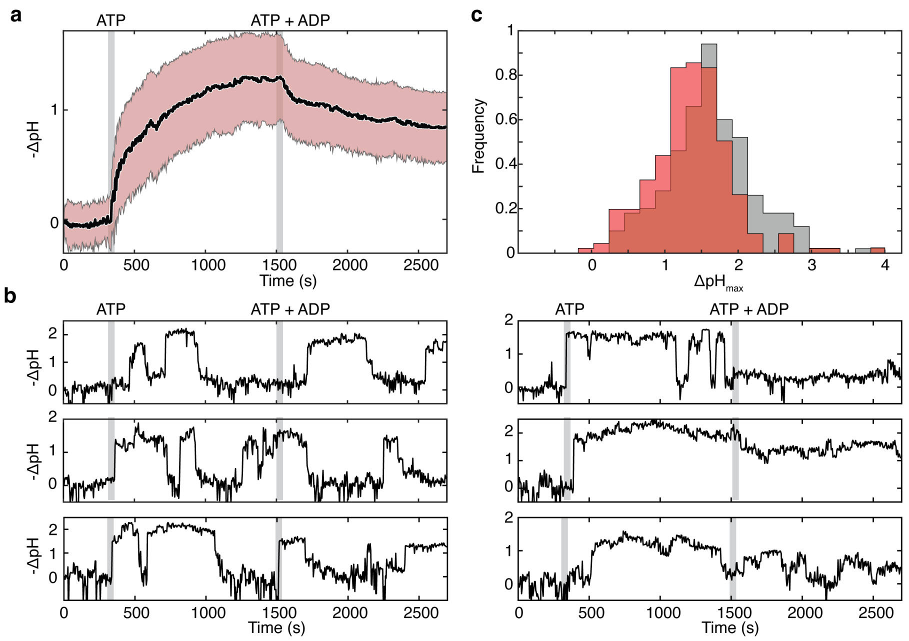


**Extended Data Fig. 14 | Additional data for ATP/ADP competition experiments.**

**a**, Ensemble average acidification kinetics. Activity was initiated upon addition of 100 μM of ATP. Subsequently, 200 μM ADP were added to the liposomal suspension. Experiments were done in the presence of 5 mM of phosphate. Black line corresponds to the mean, while shaded area corresponds to the spread of single vesicle data. **b**, Single vesicle acidification traces. **c**, Population histograms of the maximal acidification, $-\Delta pH_{max}$, of the same single vesicles in the presence of only ATP or both ATP and ADP. A two-sample Kolmogorov-Smirnov test gave a *P-value* < 0.001 (***). Total number of independent experiments was $n = 3$.

**Extended Data Table 1 | Steps involved in the sliding method.**

| |
|---|
| (1) Signal processing. Starting with $n = 5$. *Threshold* set to 0.99. |
| (2) Take first $n$ data points as the test data. Use stochastic filtering to calculate the probability $\pi_1$ of a mode-switching event in the test data, e.g. this would begin by looking for an event transitioning to an proton-pumping mode. |
| (3a) If $\pi_1 < 0.99$ (less than *threshold*): Increase the size of the test data $n = n + 5$ and return to step (2); If the end of the data set is reached in this way go to (4). |
| (3b)* If $\pi_1 \geq 0.99$ (greater than or equal to *threshold*): Take the MAP estimates for the model parameters and store the resulting estimated signal up until first event; Return to step (2) starting from the data following the MAP estimate for the event, this time looking for the alternative mode-switching events, e.g. if previously looking for proton-pumping modes, now looking for inactive modes |
| (4) When reaching the end of the data with $\pi_1 < 0.99$: Calculate the MAP estimates for the model parameters when there is no mode-switching, e.g. if the last event detected was an inactive mode; then calculate the MAP estimates when the signal remains in its inactive mode for the remaining data; Store the resulting estimated signal. |
| *In the code there is also an additional check in (3b) that there aren't two mode-switching events in the test data. This probability $\pi_2$ is calculated and if it is greater than the threshold then we use take the MAP estimates for the parameters in the model with 2 events and store the resulting estimated signal up until first event. Then going back to (2) starting from the MAP estimate for the first event. |

### References

1. Farsi, Z. *et al.* Single-vesicle imaging reveals different transport mechanisms between glutamatergic and GABAergic vesicles. *Science* **351**, 981–984 (2016).
2. Chung, S. H. & Kennedy, R. A. Forward-backward non-linear filtering technique for extracting small biological signals from noise. *J. Neurosci. Methods* **40**, 71–86 (1991).
